# A diverse landscape of FGFR alterations and co-mutations defines novel therapeutic strategies in pediatric low-grade gliomas

**DOI:** 10.1101/2024.08.27.609922

**Authors:** Eric Morin, April A. Apfelbaum, Dominik Sturm, Georges Ayoub, Jeromy DiGiacomo, Sher Bahadur, Bhavyaa Chandarana, Phoebe C. Power, Margaret M. Cusick, Dana Novikov, Robert Jones, Jayne Vogelzang, Connor C. Bossi, Seth Malinowski, John Jeang, Jared Collins, Sehee Oh, Hyesung Jeon, Amy Cameron, Patrick Rechter, Angela Deleon, Karthikeyan Murugesan, Meagan Montesion, Lee A. Albacker, Shakti H. Ramkissoon, Cornelis M. Tilburg, Emily C. Hardin, Philipp Sievers, Felix Sahm, Kee Kiat Yeo, Tom Rosenberg, Susan Chi, Karen Wright, Steve Hébert, Sydney Peck, Alberto Picca, Valérie Larouche, Samuele Renzi, Tejus Bale, Amy A. Smith, Mehdi Touat, Nada Jabado, Eric S. Fischer, Michael J. Eck, Lissa Baird, Olaf Witt, Claudia Kleinman, Quang-De Nguyen, Sanda Alexandrescu, David T.W. Jones, Keith L. Ligon, Pratiti Bandopadhayay

## Abstract

Alterations in Fibroblast growth factor receptor (FGFR)-family proteins frequently occur as oncogenes in many cancers, including a subset of pediatric gliomas. Here, we performed a genomic analysis of 11,635 gliomas across ages and found that 4.5% of all gliomas harbor FGFR alterations including structural variants (SV) and single nucleotide variants (SNV), with an incidence of almost 10% in pediatric gliomas. FGFR family members are differentially enriched by age, tumor grade, and histological subtype, with FGFR1-alterations associated with glioneuronal histologies and pediatric low-grade gliomas. Across development, we find *FGFR1* expression in both neuronal and glial precursors, while *FGFR3* expression is largely restricted to astrocytic lineages. Leveraging novel isogenic model systems, we confirm FGFR1 alterations to be sufficient to activate MAPK and mTOR signaling, drive gliomagenesis, activate neuronal transcriptional programs and exhibit sensitivity to MAPK pathway inhibitors, including pan-FGFR inhibitors. Models driven by FGFR1 SVs exhibited different patterns of sensitivity compared to those driven by SNVs. Finally, we performed a retrospective analysis of clinical responses in children diagnosed with FGFR-driven gliomas and found that targeted MAPK or FGFR-inhibition with currently available inhibitors is largely associated with stability of disease. This study provides key insights into the biology of FGFR1-altered gliomas, therapeutic strategies to target them and associated challenges that still need to be overcome.

## Introduction

Fibroblast growth factor receptor (FGFR) family proteins are commonly altered across many human cancers, including but not restricted to adult gliomas, cholangiocarcinomas, gastric, breast, lung, and bladder cancers^1–3^. The presence of FGFR alterations in these tumors has motivated the development of pan-FGFR inhibitors, of which there are four that are FDA approved for use in adult cancers. FGFR alterations have also been reported in pediatric gliomas^4–8^, however, their incidence across glioma subtypes, and optimal strategies to target them have not been fully elucidated.

Somatic driver events involving *FGFR1* have been described in pediatric low-grade gliomas (pLGG) and pediatric high-grade gliomas (pHGGs), and rearrangements in *FGFR3* were first reported in adult gliomas^4–8^. These alterations include recurrent rearrangements in FGFR1 or FGFR3, most frequently fused to TACC1 or TACC3 respectively, with FGFR1::TACC1 rearrangements occurring in pediatric gliomas and FGFR3::TACC3 in adult glioblastomas^9–11^. FGFR inhibition has been evaluated in trials of adult patients with FGFR3:TACC3 rearranged glioblastoma^12^. However, these studies were associated with limited responses, likely due to the aggressive nature of high-grade glioblastoma, as well as dose-limiting toxicities and drug resistance^2,13,14^. However, such agents have not been significantly studied in clinical trials for pediatric FGFR-driven low-grade gliomas but have potential to yield more promising results given their less aggressive nature.

pLGGs are associated with high overall survival rates, but severe life-long co-morbidities as the current standard of care treatment for children with LGGs includes surgical resection, and multi-agent chemotherapy^15^. pLGGs most commonly have driver alterations in the Mitogen-activated protein kinase (MAPK) pathway, with BRAF alterations being the most prevalent^5,6,16^. Targeted therapies including MAPK targeted therapies and BRAF inhibitor therapies have therefore been evaluated in clinical trials for patients with recurrent disease. Such efforts have yielded promising results with recent FDA approvals of the RAF inhibitor tovarafenib for recurrent BRAF-altered pLGGs, and the combination of dabrafenib/trametinib^17–19^ for pLGGs with BRAF V600E mutations. Given these drugs have primarily been directed towards patients with BRAF-altered gliomas, it is still unclear if agents such as MAPK inhibitors will be effective in the unique setting of FGFR-altered gliomas. The ideal therapeutic strategy to target these FGFR-altered pediatric gliomas remains unknown.

To begin to address this, we applied an integrative functional genomic approach to systematically evaluate the role of FGFR drivers in pediatric gliomas. We found that 4.5% of all pediatric and adult gliomas harbor FGFR alterations, with an incidence of 9.6% in pediatric gliomas. FGFR1 alterations were most frequent in pediatric gliomas, particularly pLGGs, and were sufficient to induce gliomagenesis. Generating novel isogenic neural stem cell model systems, we found FGFR1-driven model systems to be sensitive to pan-FGFR inhibition *in vitro*, with modest results observed *in vivo*. Notably, these findings are consistent with early experience in pediatric patients treated with currently available MEK or FGFR inhibition, providing key insights into the design of therapeutic approaches for children with FGFR-driven pediatric low-grade gliomas.

## Results

### Gliomas harbor a diverse landscape of alterations in FGFR1, FGFR2 and FGFR3 and are associated with specific patterns of age and co-mutations

FGFR alterations in pediatric gliomas have been described, however their incidence and association with tumor subtypes and other clinical features has not been evaluated across large cohorts of patients^4–7^. To address this, we examined a cohort of 11,635 pediatric (0-17 yrs) and adult (>18yrs) gliomas with genomic sequencing results to identify any tumors with predicted alterations in *FGFR1-4* genes. We identified 619 FGFR-altered gliomas (5.3%) across three independent datasets (DFCI, KiTZ, and Foundation Medicine) and noted that such alterations were enriched in the pediatric age group with an incidence of 9.6% (124/1288) (Supplemental Figure S1A, Supplemental Table S2) ^3,20^. Two of these datasets represented population based clinical sequencing cohorts of all patients treated with gliomas at DFCI (between 2013 and 2019) or within the Molecular Neuropathology 2.0 Study (MNP) trial led by the KiTZ^20^.

Pediatric and adult gliomas harbored recurrent but highly diverse classes of alterations in different members of the FGFR family. Alterations in *FGFR1* (50.8%, 315/619) and *FGFR3* (42.6%, 264/619) were by far the most common with *FGFR2* (5%, 31/619) and *FGFR4* (1.45%, 9/619) being altered in only a small subset of cases. *FGFR* alterations were significantly associated with age. *FGFR1* alterations were more prevalent in pediatric patients (82% of *FGFR* alterations) relative to adult patients (47% of *FGFR* alterations; fisher’s exact p<0.0001). *FGFR2* alterations were also enriched in the pediatric gliomas (12% of alterations) relative to adult gliomas (4% of alterations; fisher’s exact *p*: 0.005), while *FGFR3* alterations occurred almost exclusively in adult gliomas, also representing statistical enrichment (49% of adult *FGFR* alterations versus 6% of pediatric, fisher’s exact p<0.0001). We did not observe any significant associations between patient sex and the altered *FGFR* gene (Supplemental Figures S1B) (*p*=0.994).

FGFR-driven gliomas occur in all locations of the brain, however, different FGFR-family members exhibit anatomical predilection. Overall, 69% (81/117) of all FGFR-altered tumors in the DFCI and KiTZ cohorts were hemispheric tumors, with thalamic tumors representing the next most frequent location at 11% (13/117) (Supplemental Figure S1C). However, while FGFR2 and FGFR3 gliomas are predominantly hemispheric tumors, only ∼58 (46/79) of all FGFR1-driven gliomas were hemispheric. Instead, FGFR1 alterations also commonly drive midline gliomas that occur in the thalamus and brainstem, in addition to the cerebellum (Supplemental Figure S1C).

*FGFR1* alterations included both structural variants (SVs) (10% of FGFR-altered gliomas) and single nucleotide variants (SNVs) (40.8% of FGFR-altered gliomas) (Figure 1A, Supplemental Figure S1D-F, Supplemental Table S2), which were the most frequent. SNVs in *FGFR1* are predicted to activate the FGFR1 kinase (FGFR1 N546K and K656E mutations) (Supplemental Figure S1D) and have been previously reported in cohorts of pediatric and adult gliomas, in addition to extracranial tumors^21–25^. We also identified four other putative driver mutations of unknown significance, of which three were in the kinase domain (T141R, N546D, T657S, T658P). In contrast, alterations involving *FGFR2* and *FGFR3* were primarily structural variants (Figure 1A, Supplemental Figure S1G-H). Structural variants were most commonly gene fusion events, involving TACC1, CTNNA3 and TACC3 for FGFR1, FGFR2 and FGFR3 respectively, which have been previously reported (Supplemental Figure S1F-H, Supplemental Table S2)^9–11^. We also found 13 novel fusion partners that have not been previously reported, including PLEKHA2, the gene that encodes the protein Plekstrin Homology Domain Containing A2, which was found in two gliomas from two different cohorts (Supplemental Table S2). We also observed fusions with four fusion partners that have been previously reported in extra-cranial tumors, but not in brain tumors.

**Figure 1.**
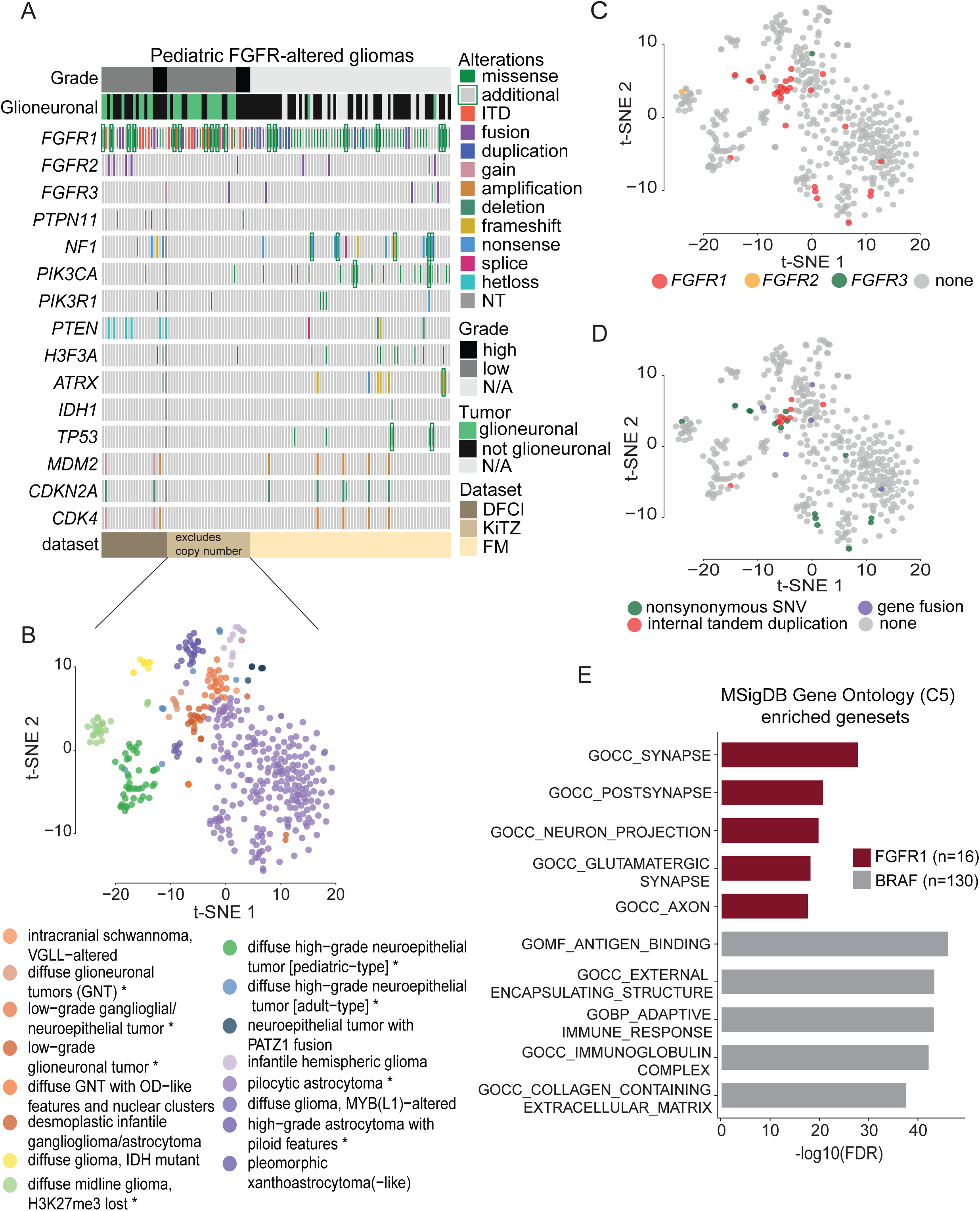
Pediatric FGFR1-altered tumors are enriched in glioneuronal histologies. A) Co-mutation plot summarizing alterations in the FGFR genes and other recurrently mutated genes across 124 FGFR-altered pediatric tumors in the three cohorts (DFCI, KiTZ, FM). Comut plot ordered by patient age, grade, and then glioma cohort. N/A-data not available. KiTZ cohort excludes copy number analysis. B) Unsuper-vised nonlinear t-distributed stochastic neighbor embedding (t-SNE) of DNA methylation profiles of 425 pediatric gliomas from the KiTZ cohort. Each sample is color-coded in their respective tumor class color. Tumor class names with an asterisk (*) represents histologies with FGFR1 alterations. TSNE plots of the 425 tumors colored by C) the specific FGFR gene altered or D) the FGFR alteration type. E) Horizontal bar plots depicting the top 5 significant Gene Ontology C5 (MsigDB) terms enriched (ranked by significance) in FGFR1 (n=16) or BRAF-altered (n=130) gliomas. Negative log(FDR) values from the GSEA analysis are shown on the x-axis. FM: Foundation Medicine.

These include AMBRA1, OPTIN, SHTN1 and GKAP1, the genes that encode the proteins Autophagy and Beclin 1 Regulator 1, Optineurin, Shootin 1, G Kinase Anchoring Protein 1, respectively (Supplemental Table S2). The second most common FGFR1 structural variant observed was an internal tandem duplication (ITD) of the FGFR1 kinase domain, which occurred at a similar incidence to prior reports (Figure 1A, Supplemental Figure S1E)^4–6^. We observed novel recurrent *FGFR4* alterations (V508M, R80W, D425N, V550I, and amplifications) but these were rare occurring only in nine adult patients within the Foundation Medicine cohort. *FGFR1-ITD* and *FGFR2* SVs were only found in pediatric gliomas, whereas *FGFR1/3* gain, *FGFR3* fusion, and *FGFR4* alterations were almost exclusively found in adult gliomas (Supplemental Figure S1I).

We also observed significant associations between the different FGFR-family members and histological features of the gliomas, with FGFR1 alterations found most frequently in glioneuronal tumors. FGFR1-driven gliomas spanned both pediatric low-grade and high-grade gliomas (Figure 1A, Supplemental Figure S1J). While both SNVs involving FGFR1 occurred in both low-grade and high-grade gliomas, the FGFR1 structural variants were most frequent in pediatric low-grade gliomas. In contrast, gliomas with FGFR2 SVs were exclusively observed in pediatric low-grade gliomas while FGFR3-rearranged gliomas were primarily observed in adult high-grade gliomas (glioblastoma IDH wild-type) (Supplemental Figure S1J). Diagnostic classification based on DNA methylation profiling was available for 425 gliomas within the KiTZ dataset (Figure 1B-D). Integrating DNA methylation profiles with matched DNA-sequencing, we observed *FGFR1* alterations in 5.4% of pediatric low-grade gliomas (23/425), with 70% (16/23) of these tumors among the glioneuronal subtypes (Figure 1B-D). We similarly found 50% (9/18) of pLGGs with *FGFR1* alterations were morphologically classified as glioneuronal tumors in the DFCI cohort (Figure 1A).

One of the most distinctive aspects of pLGGs with *FGFR1* SNVs was that they frequently co-occurred with other driver events. This is in contrast with BRAF-altered or pLGGs driven by other alterations, which are thought to largely represent single-driver tumors ^4,26^. Within pLGGs in the DFCI and KiTZ cohorts, 80% (16/20) of all gliomas with *FGFR1* SNV harbored at least one additional somatic event, either involving *FGFR1* itself, or additional driver alterations predicted to activate mTOR or MAPK signaling (*NF1* (n=2), *PTPN11* (n =2), *PIK3CA* (n=1), *PIK3R1* (n=1) (Figure 1A, Supplemental Table S2)^4–6,16,22,27^. These consisted of pathogenic SNVs in *PTPN11* including G503R and E76Q SNVs that are in located in the phosphatase domain and the SH2 domain of the protein, respectively. In contrast, only 10.5% (2/19) of pLGGs driven by *FGFR1* SVs harbored co-occurring alterations. *FGFR1* SNVs in pediatric HGGs also co-occurred with other driver events that have been described in pediatric HGGs, however, these differed from those observed in pLGGs and included histone mutations, copy-number alterations including *MDM4* and *CDK4* amplifications, and deletions in *CDKN2A/B* (Figure 1A, Supplemental Table S2).^7,8,16^

In addition to the presence of *FGFR1* alterations co-occurring with mutations in genes in the MAPK or PI3K pathway, they most frequently co-occurred with additional mutations in *FGFR1* itself in 55% (11/20) of pLGGs. We found that ∼27% (3/11) of patients had a co-occurring mutation that occurred in the extracellular domain (A21V, G35R, R148S), which all occurred concurrently with the N546K hotspot mutation within the FGFR1 kinase (Supplemental Figure S1D). The remaining eight patients had co-occurring mutations in the kinase domain with ∼54.5% (6/11) harboring co-occurring mutations with only the K656E mutation (G660C, D683G, D652G, K638R, H649R, N546S, K655E, I651M—some tumors have multiple co-occurrences), while ∼18% (2/11) co-occurred with both the K656E and N546K mutations (V561M, K656N) (Supplemental Figure S1D).

Altogether, these data provide evidence that FGFR alterations are frequent in ∼10% of pediatric gliomas and these tumors have heterogeneous FGFR alterations patterns with both SNV and SVs. In addition, FGFR1-altered gliomas were more enriched in glioneuronal tumor types in pediatric patients.

### Expression profiles of FGFR gliomas identifies neuronal signatures and lineage associations to normal stages of brain development

Our observation that FGFR1-altered gliomas are predominantly classified as glioneuronal tumors by DNA methylation profiling raises the possibility that they may also express neuronal differentiation and signaling pathway genes. To investigate this, we interrogated transcriptome-wide expression profiles of 146 pediatric gliomas, 16 FGFR1-driven (9 SVs and 7 SNVs) and 130 BRAF-driven tumors (88 SVs, 42 SNVs)^28^ to identify gene signatures that are differentially expressed in FGFR-altered gliomas (Supplemental Table S3). Indeed, gene set enrichment analysis (Gene Ontology C5 gene sets in MsigDB)^29,30^ of the 206 genes significantly upregulated in FGFR1-altered tumors revealed enrichment of neuronal gene sets specifically involving the synapse and neurotransmission (Figure 1E, Supplemental Table S3). In addition, within the Cell Type signatures (C8 gene sets in MsigDB)^29,30^, gene sets enriched within the FGFR1-driven gliomas were predominantly represented by multiple primary neuronal cell types, including GABAergic neurons and dopaminergic neurons (Supplemental Figure S2A). In addition, GSEA analysis including gene sets within the C2 curated database revealed enrichment of signatures of polycomb repression and promoter bivalency in stem and progenitor cells (Supplemental Figure S2B), which have been associated with regulating expression cell differentiation programs. In contrast, BRAF-altered gliomas showed enrichment of gene sets that regulate immune response and extracellular matrix/matrisome pathways (Figure 1E, Supplemental Figure S2B), including those involved with differentiation of stromal and fibroblast cell types^31^ (Supplemental Figure S2A). These data provide evidence that FGFR1-altered gliomas are associated with clear enrichment of neuronal expression signatures compared to their BRAF-altered counterparts. This differential lineage enrichment between FGFR1 and BRAF-altered tumors may be reflective of developmental origins of the tumors, tumor location, or the consequence of different driver mutations.

The striking association between different FGFR family drivers across different classes of pediatric and adult gliomas led us to hypothesize that these differences reflect differences in the developmental origins of each glioma. We tested this by investigating temporal and spatial expression of the FGFRs in prenatal development through postnatal life, across brain regions and cell types. We first profiled expression of each FGFR family member across time (prenatal to adult brain) and brain regions, using two human lifespan datasets of bulk RNA-sequencing, the Evo-devo atlas^32^ and Brainspan^33^ (Figure 2A, Supplemental Figure S3A-D).

**Figure 2.**
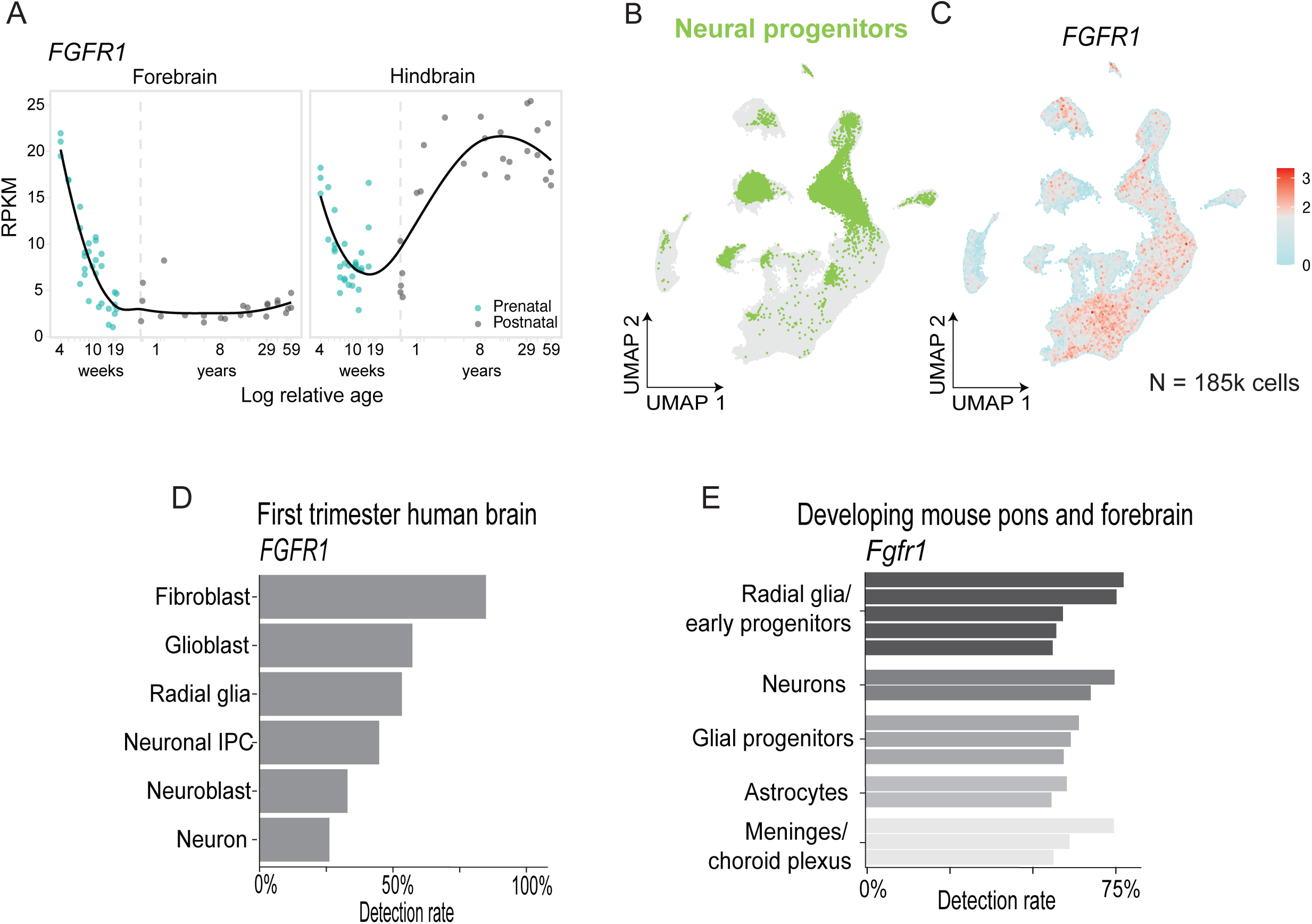
Expression of FGFR1 is found in neuronal and glial lineage cell types in brain development. A) Expression of FGFR1 in bulk RNA-seq across lifespan in human forebrain (n=55) (left) and hindbrain (n=59) (right). X-axis denotes sample age, measured in natural log of weeks post conception, and x-axis tick values correspond to weeks in prenatal points and years in post-natal points. Vertical dashed line represents birth. Y-axis depicts RPKM expression values. B-C) UMAP representation of human embryo cells 4-5.5 weeks post conception. B) Left: Cells colored by neural progenitor cells annotated in the original study colored green. C) Right: Cells colored by normalized expression of FGFR1. Color scale midpoint is set to the midpoint of maximum and minimum expression. D) Detection rate (% of cells with expression >0) of FGFR1 in broad cell types of first trimester human brain. E) Top 15 cell clusters by detection rate (% of cells with expression of >0) of Fgfr1 in developing mouse pons and forebrain. Cell clusters grouped by broad cell type.

Expression of both *FGFR1* and *FGFR3* exhibits temporal association with development and are both expressed in early development. In the forebrain, *FGFR1* is expressed early in the prenatal brain and decreases towards birth (Figure 2A, Supplemental Figure S3A). In the cerebellum, in contrast, while prenatal patterns of *FGFR1* expression mirror those of the forebrain, postnatal expression continues to increase through life, a result corroborated across datasets^34^ (Figure 2A, Supplemental Figure S3A). In turn, *FGFR3* displayed similar patterns for the forebrain and hindbrain, with higher expression in the first weeks of prenatal development (Supplemental Figure S3D). FGFR2 showed very low expression with minimal changes over developmental time (Supplemental Figure S3C).

We next hypothesized that the association of FGFR1-alterations with glioneuronal gliomas suggests that these gliomas may arise from a specific types of undifferentiated neural progenitor cells, thereby giving rise to multipotent glioneuronal tumors. We therefore assessed *FGFR1* expression in single-cell RNA expression atlases of the human embryo 4-5.5 pcw^35^, human brain in the first trimester of gestation^36^, as well as murine atlases of the developing forebrain and pons^37,38^. Indeed, we found that *FGFR1* expression was not restricted to any one cluster or cell type in the brain and instead was observed across multiple lineages, including glial cells, neuronal cells, neural and glial progenitors, and non-neuroectoderm cells across both human and mouse datasets (Figure 2B-E, Supplemental Figure S3G).

Finally, we evaluated associations between expression of *FGFR2* and *FGFR3*, and normal cells across development. *FGFR2* was expressed highly in the spinal cord (Supplemental Figure S3E) and enriched within multiple cell types including the choroid plexus cells, oligodendrocytes, and astrocytes both pre- and postnatally in single-cell mouse forebrain and pons (Supplemental Figure S3F). *FGFR3* is more abundantly expressed than *FGFR1* and *FGFR2* in bulk data but does not appear to demonstrate lineage specificity. Very early in development, before initiation of gliogenesis in humans and correlating with the prenatal peak observed in bulk datasets, *FGFR3* is expressed in neural progenitors (Figure 2B, Supplemental Figure S3G, 3H). Later on, *FGFR3* is most strongly detected in astrocytes, across regions and across species, with expression sustained in this lineage in the adult brain (Supplemental Figure S3I-J). This is intriguing as FGFR3 alterations are predominantly found within glioblastoma in adults.

Overall, the presence of *FGFR1* expression in both glial and neuronal cell types in the developing brain may suggest its expression is associated with early progenitor cells with similarity to phenotypes seen in glioneuronal tumors where they occur.

### Expression of FGFR1 alterations is sufficient to induce growth factor independence

Together, our data reveals FGFR alterations as prevalent recurrent drivers across gliomas and highlight the specific association between FGFR1 alterations and pediatric gliomas. However, the mechanisms through which these events drive pediatric glioma growth, including cooperation with co-occurring events, and strategies to target them have not been fully determined.

To corroborate FGFR1 as a potential therapeutic target in pLGGs, we sought to evaluate its role in driving gliomagenesis. There are no patient-derived FGFR1-pLGG models as the tumors cannot be propagated *ex vivo*. Therefore, we generated a panel of isogenic primary mouse neural stem cells (mNSC) and Tert-immortalized human neural stem cells (ihNSCs) models that were transduced to express FGFR1 and other driver alterations that are found in pLGGs (Figure 3A). In total we generated a comprehensive collection of nine novel lines including FGFR1 N546K (F1-N546K), FGFR1-ITD (F1-ITD), FGFR1::TACC1 (F1::TACC1), PTPN11 SNV (PTPN11 alone) as well as a line with the double mutations in FGFR1 and PTPN11 (F1-N546K+PTPN11). These were compared to lines with wild-type FGFR1 (F1-WT), BRAF drivers common in pLGGs (KIAA1549::BRAF fusion or BRAF V600E SNV) and GFP or HcRed (Vector control) as the control (Figure 3A, Supplemental Figure S4A-B).

**Figure 3.**
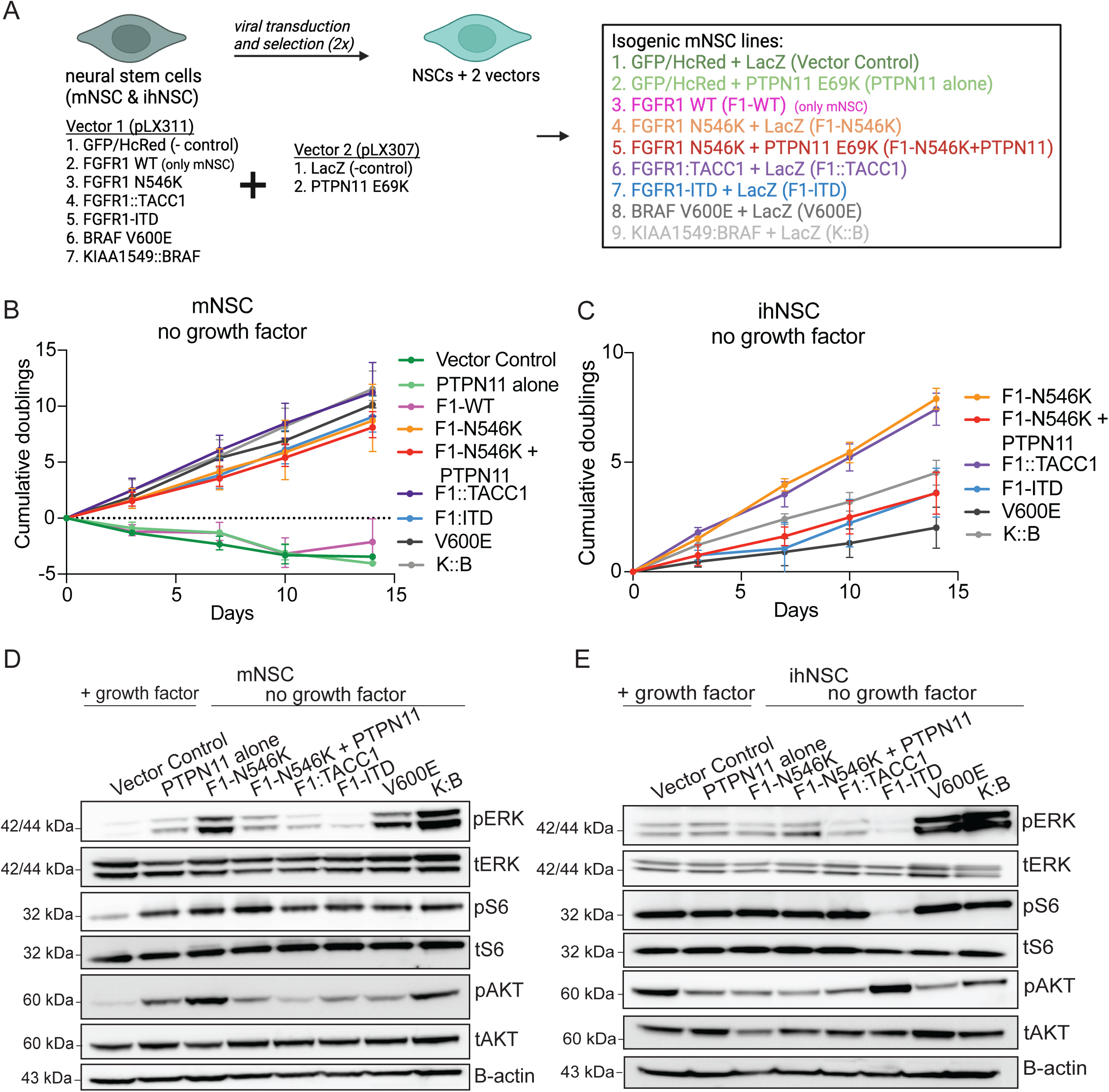
Isogenic NSC models driven by FGFR1 alterations grow independently of growth factor. A) Overview of the generation of the isogenic NSC lines. Mouse or Tert-immortalized human NSCs were virally transduced with each of the FGFR or BRAF alterations, or control vectors, followed by a second transduction containing the co-occurring PTPN11 mutation or control vectors. Abbreviations for models= F1-N546K: FGFR1 N546K SNV, F1-N546K + PTPN11: FGFR1 N546K SNV + PTPN11 E69K SNV, F1-ITD: FGFR1-ITD, F1::TACC1: FGFR1::TACC1, V600E: BRAF V600E SNV, K::B: KIAA1549::BRAF. B) Cumulative doubling growth curves for the isogenic mNSC lines in the absence of exogenous growth factor. Values and error bars represent the average +/- SEM of three independent experiments. C) Cumulative doubling growth curves for the isogenic ihNSC lines in the absence of exogenous growth factor. Values and error bars represent the average +/- SEM of three independent experiments. D) Represent western immunoblots of downstream MAPK and PI3K/mTOR signaling pathways effectors. Phosphorylated proteins represent activated signaling pathways. pERK is a readout for MAPK signaling and pAKT and pS6 are readouts for PI3K/mTOR signaling. Vector Control and PTPN11 alone are grown in the presence of growth factor, while lines harboring FGFR1 and BRAF drivers are grown without growth factor supplementation.

We first evaluated whether expression of FGFR1 alterations were sufficient to render our mNSCs and ihNSCs growth factor independent, a hallmark of transformation. Wild-type mNSCs and ihNSCs require supplementation of growth factors including epidermal growth factor (EGF) and fibroblast growth factor (FGF2) to maintain stemness and proliferative potential. We leveraged this model system to withdraw EGF and FGF2 from each of our engineered cell lines and measured cell proliferation in the presence and absence of growth factors. All murine and human models grew robustly in the presence of growth factors (Supplemental S4D-F). mNSC models driven by FGFR1 alterations were able to maintain proliferation in the absence of growth factors and showed similar growth to BRAF positive controls (Figure 3B). No growth was seen in FGFR1 WT, PTPN11 SNV, or vector control. We also observed similar trends in the ihNSC where all FGFR1 alterations are sufficient to induce growth-factor independence which was similar to BRAF while cells transduced with PTPN11 SNV or vector control exhibited slow growth in the absence of growth factor (Figure 3C and Supplemental Figure S4E-F).

We next evaluated whether the FGFR1 alterations are sufficient to maintain downstream MAPK and mTOR signaling to bypass the need for growth factor supplementation. Leveraging immunoblots to assess levels of total and phosphorylated effector kinases, we observed activation of both pathways across our panels of FGFR1 and BRAF-driven NSCs, probing ERK, S6 and AKT, and their phosphorylated, and thus, activated forms (Figure 3D-E, Supplemental Figure S4G-L). Importantly, the level of activation observed in FGFR1-altered models in the absence of growth factor was similar to that observed with vector control NSC lines in the presence of growth factors (Figure 3D-E, Supplemental Figure 4G-L). We observed a trend of higher ERK activation in the BRAF-altered models, however, this did not reach statistical significance.

These data lead us to conclude that diverse FGFR1 alterations are each sufficient to activate MAPK and mTOR signaling and to induce growth factor independence in mouse and human NSC models.

### FGFR1-altered mNSC models are enriched in neuronal transcriptional programs

We next sought to determine whether our isogenic mNSC FGFR1 model systems transcriptionally resemble human FGFR1-driven gliomas. We performed bulk RNA-sequencing on our mNSC models transduced to express the FGFR1 or BRAF alterations. FGFR1-altered lines were more similar to each other by PCA and correlation analysis than to the BRAF and vector control line (Supplemental Figure S5A-B). We found a total of 659 genes to be differentially expressed between the FGFR1 and BRAF-altered lines (FDR < 0.05, absolute LFC 1.5) (Figure 4A, Supplemental Table S4). Of these, 319 genes were upregulated in the FGFR1-altered models. The most differentially expressed gene was *Hs6sp2*, that encodes Heparin Sulfate 6-O Sulfotransferase 2 and is important for efficient signaling of FGFR family receptors^39^. Multiple genes involved in neuronal development including *Epha3*, *Emb*, *Ednrb* and *Ntrk2* were also within the top 25 most differentially expressed genes in FGFR1 models (Figure 4A).

**Figure 4.**
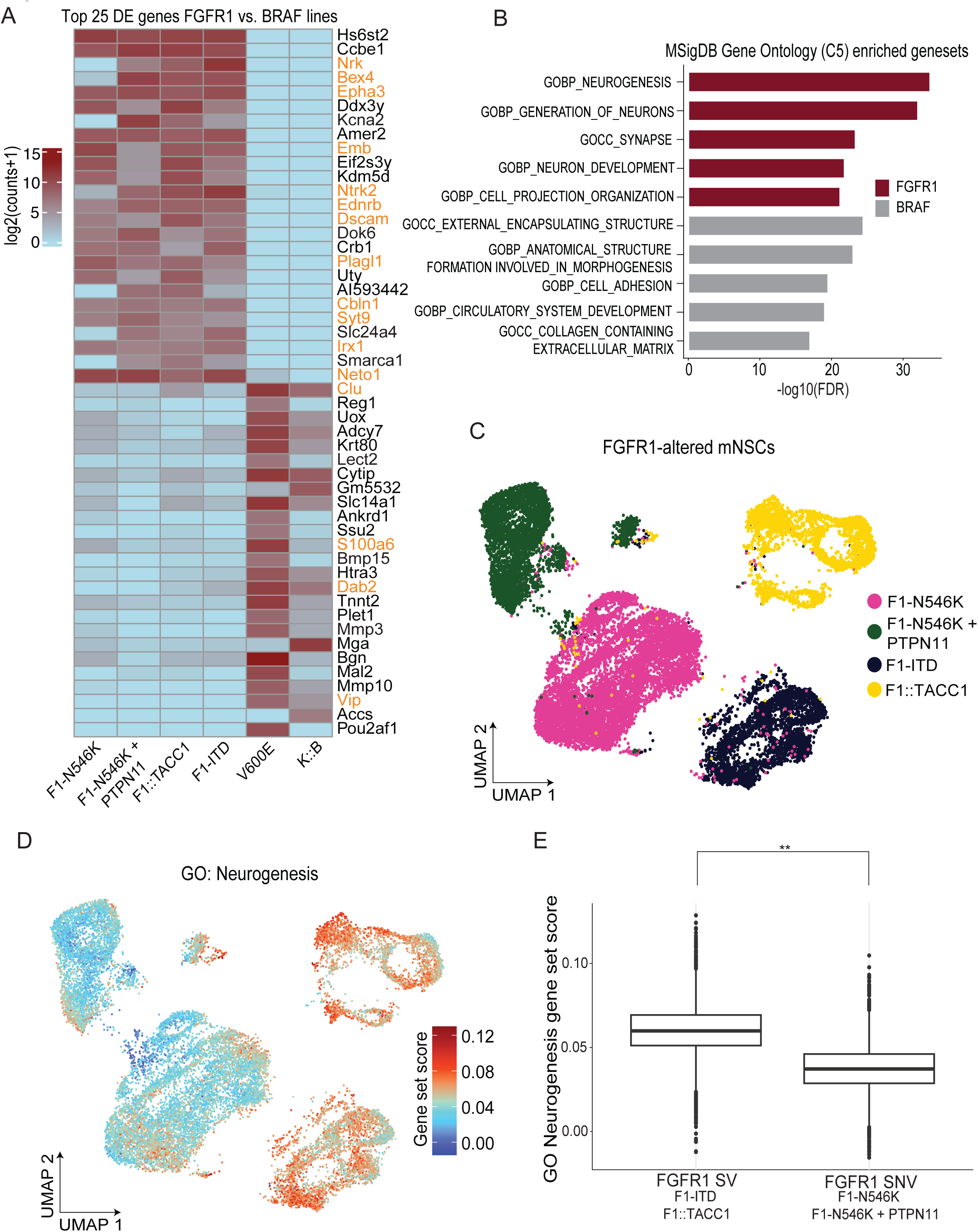
FGFR1-altered lines are enriched in similar neuronal signatures observed in patient tumors. A) Heatmap of the top 25 genes differentially expressed in the FGFR1 vs BRAF-altered lines. Gene names highlighted in orange represent genes involved in neuronal development. B) Horizontal bar plots depicting the top 5 significant Gene Ontology C5 (MsigDB) Terms enriched (ranked by significance) in FGFR1 (n=4) or BRAF (n=2)-altered mNSC lines. C) UMAP embedding of all FGFR1-altered mNSC lines, colored by line. D) UMAP embeddings of the FGFR1-altered mNSCs colored by the GO Neurogenesis gene set. Legend depicts gene set score. E) Box plots quantifying the expression of the GO Neurogenesis gene set score in the FGFR1 SV and SNV samples. **p<0.0001, Wilcoxon Rank Sum test. Abbreviations for models= F1-N546K: FGFR1 N546K SNV, F1-N546K + PTPN11: FGFR1 N546K SNV + PTPN11 E69K SNV, F1-ITD: FGFR1-ITD, F1::TACC1: FGFR1::TACC1, V600E: BRAF V600E SNV, K::B: KIAA1549::BRAF.

Indeed, applying gene set enrichment analysis, we found that expression programs within NSCs transduced with the FGFR1 oncogenes were significantly enriched with gene sets associated with neuronal differentiation when compared to BRAF-expressing cells, similar to our prior analyses across the human pLGG samples (Figure 4B, Supplemental Figure S5C-D, Supplemental Table S4). In addition to neuronal program gene sets, we also found enrichment of gene sets associated with neuronal cell signatures, and polycomb repressive marked genes (Supplemental Figure S5C-D). In contrast, within the BRAF-altered cell lines we saw enrichment of extracellular matrix (ECM) programs, adhesion and motility terms, and stromal and endothelial cell signatures (Figure 4B, Supplemental Figure S5C-D, Supplemental Table S4), also similar to our prior findings in BRAF-altered patient gliomas (Figure 1E, Supplemental Figure S2A-B).

This expression of neuronal expression programs within the FGFR1 models is heterogenous. We evaluated the heterogeneity of transcriptional signatures identified in the FGFR1-altered cell lines and patient tumors (Figure 1E) using single-cell RNA-sequencing of the FGFR1-altered mNSC lines (Figure 4C). Using Uniform Manifold Approximation and Projection (UMAP) embedding for dimension reduction analysis, we found that the different FGFR1-altered mNSC lines clustered separately from each other (Figure 4C), independent from expression of cell-cycle signatures within individual cells (Supplemental Figure S5E). FGFR1-altered cells exhibited heterogenous expression of the GO neurogenesis gene set, both within and across cell lines (Figure 4D). The FGFR1 SV-driven lines had significantly higher expression of the GO neurogenesis gene set compared to the FGFR1 SNV-driven lines, suggesting different FGFR1 alterations may influence these neuronal transcriptional states differently (Figure 4E).

These data suggest that our isogenic FGFR1-mutant mNSCs express similar cell programs as the patient FGFR1-altered gliomas, with enrichment of gene sets associated with neuronal differentiation, development and signaling compared to BRAF-driven glioma models.

### FGFR1 alterations are sufficient to drive tumor formation *in vivo*

Having confirmed that our models recapitulated expression programs observed in human pLGGs, we next sought to evaluate whether each FGFR1-alteration was sufficient to induce gliomagenesis. We orthotopically injected each isogenic mNSC models into the brains of SCID mice and evaluated glioma formation (Supplemental Figure S6A). Mice harboring the vector control or PTPN11 E69K alone cells did not form any tumors and there was one non-tumor related death (Figure 5A-B). Intracranial injection of mNSC transduced to express our positive controls (BRAF V600E and KIAA1549::BRAF) was sufficient to induce glioma formation, with BRAF V600E resulting in more consistent and rapid tumor formation than the KIAA1549::BRAF fusion and FGFR1 alterations (Figure 5B).

**Figure 5.**
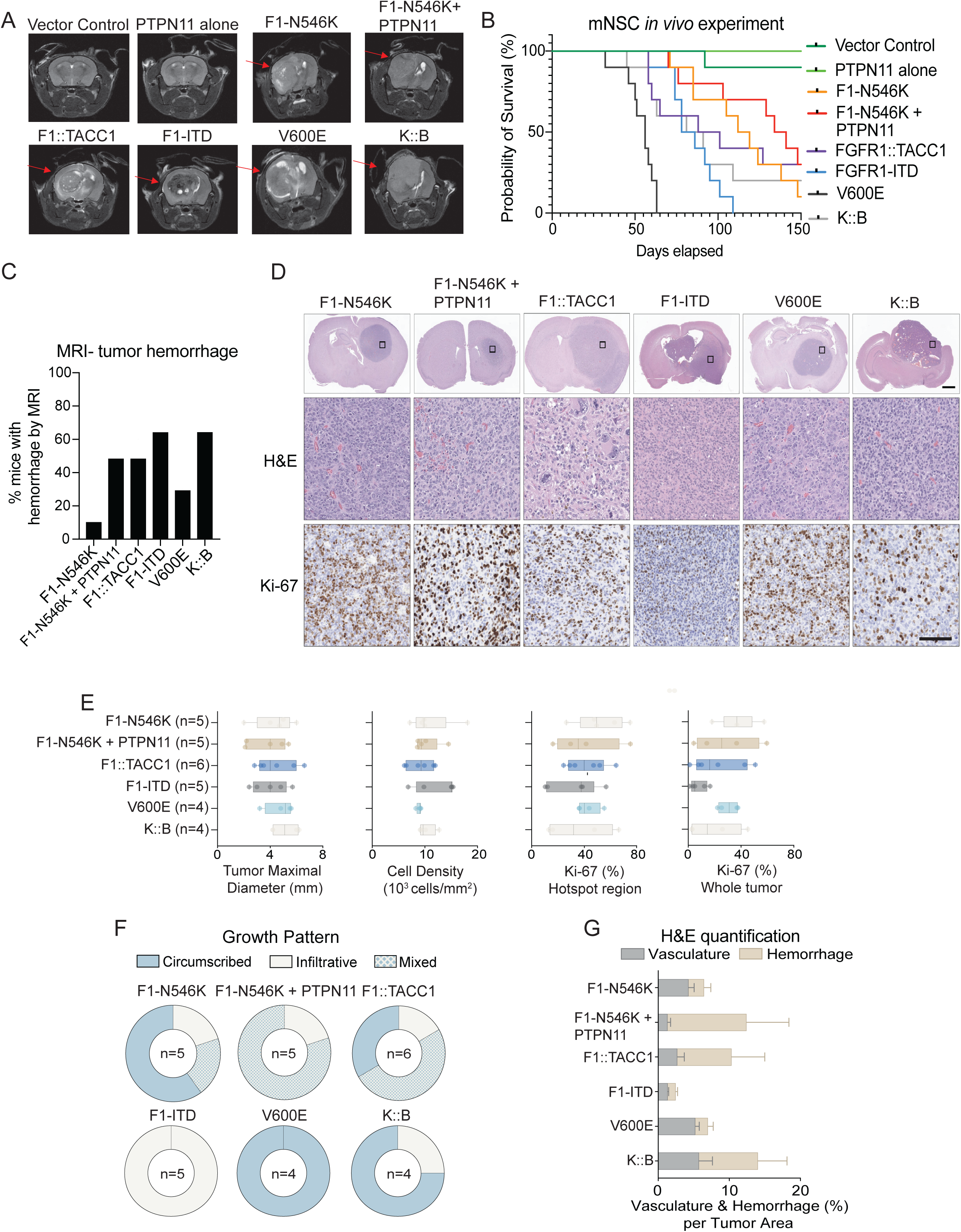
Isogenic mNSC lines driven by FGFR1 alterations form tumors in mice. A) Representative axial MRI images of gliomas form following intracranial injection of the isogenic mNSC lines expressing oncogenes shown. Abbreviations for models= F1-N546K: FGFR1 N546K SNV, F1-N546K + PTPN1: FGFR1 N546K SNV + PTPN11, F1-ITD: FGFR1-ITD, F1::TACC1: FGFR1::TACC1, V600E: BRAF V600E, K::B: KIAA1549::BRAF. Red arrow points to gliomas in the right hemisphere of the brain. B) Kaplan-Meier survival curves of mice harboring intracranial allografts of mNSCs transduced to express each alteration. Ten mice were injected with each cell line. C) Bar graph showing the percentage of mice with tumor hemorrhage by MRI assessment. # of mice is depicted. D) Representative H&E images showing glioma formation at low magnification (1.5x, scale bar = 1 mm) and high magnification (40x, scale bar = 100 μm), with H&E slides and Ki-67 staining. E) AI-based digital quantification of tumor maximal diameter, cellularity, and Ki-67 positivity in hotspot regions and whole tumors using U-net on the Visiopharm platform. See Methods for detailed quantification schema. # of mice is depicted. F) Distribution of tumor border’s growth pattern (infiltrative, circumscribed, or mixed) for each group. # of mice is depicted. G) AI-based digital quantification of vasculature and hemorrhage percentage per tumor area in each group using DenseNet on the HALO AI platform. Data presented as mean ± SEM.

MRI imaging was used to track glioma formation during this study (Figure 5A, Supplemental Figure S6A). A total of 78 mice (58 with FGFR1 or BRAF alterations) were imaged during the study, and gliomas were confidently detected in ∼72% mice implanted with mNSCs driven by FGFR1 or BRAF alterations. The overall tumor penetrance for BRAF V600E transduced mNSCs was 100%, while the penetrance of KIAA1549::BRAF was 90%. Injection of mNSCs transduced to express either of the FGFR1 structural variants (FGFR1::TACC1 or FGFR1-ITD) was also sufficient to induce glioma formation with an overall penetrance of 100%. The overall penetrance of tumor formation following injection of NSCs transduced to express the FGFR1 N546K mutation alone was 80%, while the penetrance for NSCs transduced to express both the FGFR1 N546K and PTPN11 mutation was 90%.

Intracranial hemorrhage was noted in ∼45% of all gliomas and occurred least frequently in gliomas driven by BRAF V600E or by FGFR1 N546K alone. Within the FGFR1-altered gliomas that underwent MRI (n=39) intracranial hemorrhage was detected in 50% (5/10) of gliomas driven by FGFR1 N546K + PTPN11E69K, 50% (5/10) in FGFR1::TACC1-driven gliomas, and 66.66% (6/9) in FGFR1-ITD driven gliomas (Figure 5C). In contrast, only 10% (1/10) of the FGFR1 N546K gliomas had intracranial hemorrhage. Within the BRAF-altered gliomas that underwent MRI (n=19), intracranial hemorrhage was detected in 30% (3/10) of BRAF V600E tumors and 66.66% (6/9) in gliomas that harbor the KIAA1549::BRAF fusion (Figure 5C).

We observed differences in overall survival of mice following intracranial injection of each of these isogenic model systems. Mice injected with BRAF V600E-expressing NSCs had the shortest overall survival compared to all other conditions (median survival of 56 days, *p* value <0.001), while mice harboring allografts of the mutant FGFR1 N546K+PTPN11 mutant NSCs survived the longest (median survival of 141 days) (Figure 5B). Interestingly, mice harboring allografts transduced to express the FGFR1 SVs (FGFR1-ITD and FGFR1::TACC1) exhibited a trend to shorter survival compared to those with the FGFR1 SNVs (82 and 94.5 days respectively, compared to 141 for the double mutant and 115.5 days for the FGFR1 N546K alone) (Figure 5B).

Neuropathological and AI-aided examination of the tumor histology and immunostaining showed mNSC FGFR1 and BRAF-altered tumors all had pathological features consistent with a glioma (Figure 5D, Supplemental Figure S6B & S6D-E). Cells had glial cytology in many regions, but all showed mitotic activity and moderate to severe atypia most consistent with higher grade, not lower grade, gliomas. No specific features of low-grade gliomas were noted (Rosenthal fibers, eosinophilic granular bodies, biphasic appearance, ganglionic neurons). The tumors showed glial lineage expression and AI-aided quantification of glial makers (GFAP and OLIG2) and Ki67 as a marker of proliferation showed differences in their patterns of expression (Figure 5E, Supplemental Figure S6C). All genotypes expressed GFAP and OLIG2 and the average Ki67 proliferation rate across all tumors was 24% but with a wide range in growth rates across genotypes (1.13% - 59.5%) (Figure 5D-E, Supplemental Figure S6B-C). We did not observe significant differences in tumor diameter, cell density Ki67 proliferation rate, or glial marker staining between different FGFR1-genotypes (Figure 5E, Supplemental Figure S6C).

The tumors did show strikingly different levels of diffuse infiltration of the brain parenchyma and other areas of assessment of tumor diameter, density, using AI-aided classifiers we previously trained on human gliomas (HALO-AI) (Supplemental Figure S6F). FGFR1-ITD gliomas were scored as diffuse and did not have areas of circumscribed tumor. In contrast, BRAF V600E gliomas were scored as completely circumscribed with pushing borders. The other glioma genotypes all harbored mixed growth patterns (Figure 5F).

Given our observations of intracranial hemorrhage detected by MRI imaging, we also evaluated the incidence of hemorrhage within these samples by assessing the percentage of hemorrhage relative to vasculature within each glioma also using our AI-aided classifier for features of glioma (Supplemental Figure S6G). Gliomas expressing KIAA1549:BRAF, FGFR1:TACC1 and FGFR1 N546K + PTPN11 had the highest proportion of hemorrhage relative to vasculature, while those with BRAF V600E gliomas had the lowest (Figure 5G).

### pan-FGFR inhibitors represent therapeutic opportunity for FGFR1-driven gliomas

Small molecule inhibitors targeting the MAPK and mTOR pathways have rapidly emerged as novel therapeutic approaches for children with pLGGs, including recent FDA approvals for the use of pan-RAF inhibitors, or combination MEK and Type 1 BRAF inhibition, for gliomas that harbor BRAF alterations^17,18,40^. However, the activity of these agents in FGFR1-driven tumors is unknown, and the most efficacious path to precision medicine approaches for children diagnosed with FGFR1-driven pLGGs is unclear. We thus evaluated these agents across our isogenic models.

We first evaluated the *in vitro* efficacy of a panel of MEK (trametinib), RAF (belvarafenib or tovorafenib), and mTOR (everolimus) inhibitors across our panel of isogenic mouse and human NSCs (Supplemental Figure S7A). We found that our oncogenic BRAF mNSC models exhibited sensitivity to trametinib and the RAF inhibitors (Figure 6A-C, Supplemental Figure S7B) with minimal activity observed with single agent everolimus (Figure 6D, Supplemental Figure S7C). Focusing more closely on the MEK and RAF inhibitors, we found that the FGFR1-driven NSCs were less sensitive to these agents compared to the BRAF-driven models (Figure 6A-C, Supplemental Figure S7B). However, we did also observe some variability in sensitivity within the FGFR1-driven lines, with models harboring FGFR1 SNVs exhibiting greater sensitivity than the FGFR1 SVs (Figure 6A-C, Supplemental Figure S7A-B).

**Figure 6.**
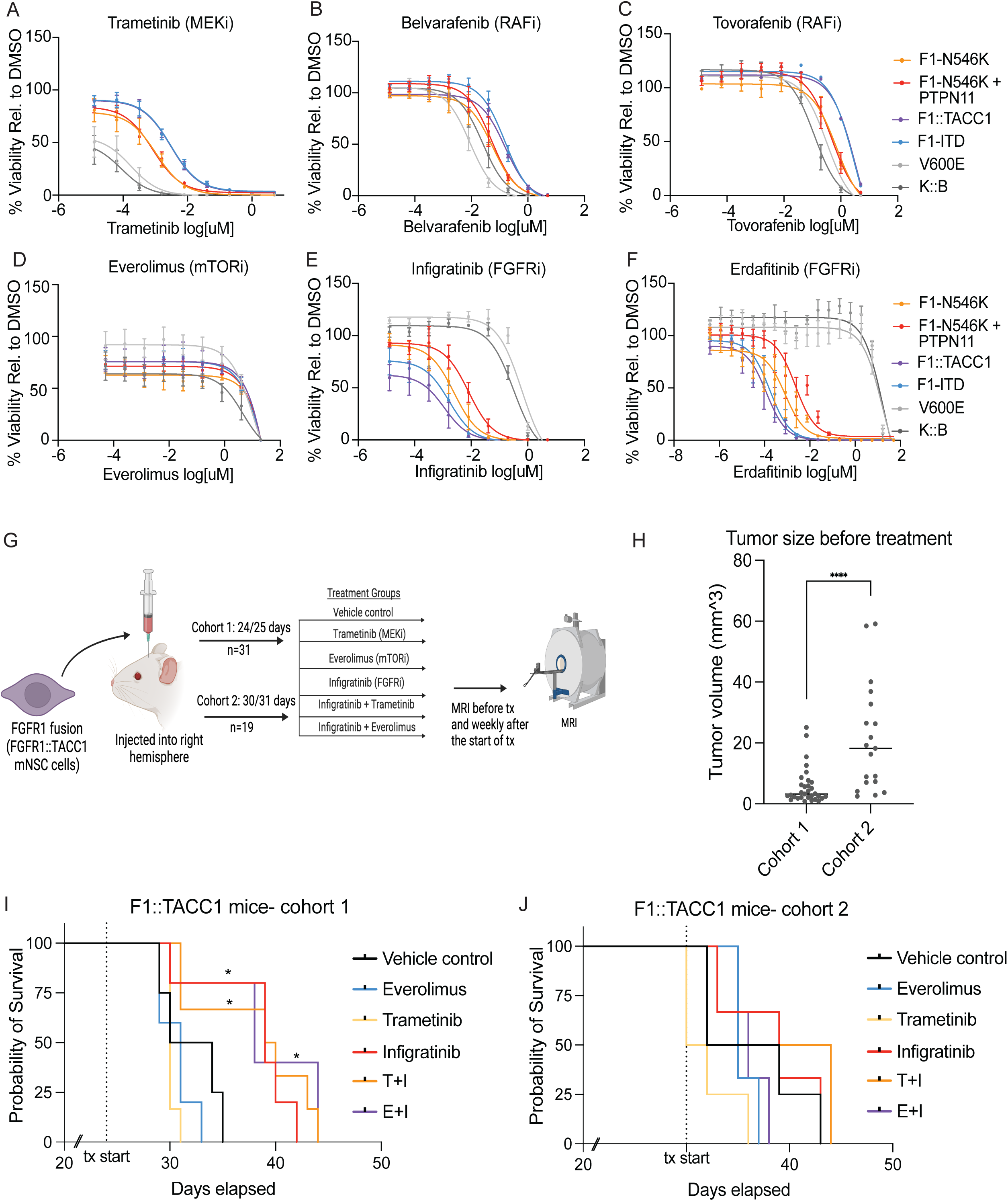
FGFR1-driven models are sensitive to FGFR inhibition. Dose response curves in the FGFR1 and BRAF-altered mNSC models for A) trametinib (MEKi), B) belvarafanib (RAFi), C) tovorafenib (RAFi), D) everolimus (mTORi), E) infigratinib (FGFRi), F) erdafitinib (FGFRi). G) Model summarizes the in vivo drug study. 300,000 F1::TACC1-driven mNSC cells were injected into the right hemisphere of mice. After gliomas were detected, mice were treated with single or combinations of infigratinib, everolimus, and trametinib. MRI was performed prior to drug treatment and weekly after start of drug treatment. Cohort 1 (n=31) were treated after 24 or 25 days post cell injections and Cohort 2 (n=19) were treated after 30 or 31 days post cell injections. H) Dot plot showing the volume of tumors in each cohort prior to treatment. Volume assessed by MRI. **** p<0.0001. I) Kaplan-meier survival curves for mice in Cohort 1 by each treatment group (treated after 24/25 days-dotted line). J) Kaplan-meier survival curves for mice in Cohort 2 by each treatment group (treated after 30/31 days-dotted line). * p<0.05. Abbreviations for models= F1-N546K: FGFR1 N546K SNV, F1-N546K + PTPN11: FGFR1 N546K SNV + PTPN11 E69K SNV, F1-ITD: FGFR1-ITD, F1::TACC1: FGFR1::TACC1, V600E: BRAF V600E SNV, K::B: KIAA1549::BRAF.

In contrast, all FGFR1-altered models were more sensitive to pan-FGFR inhibitors. We performed dose response curves in our FGFR1 and BRAF-altered lines using four FDA-approved pan-FGFR inhibitors (infigratinib, erdafitinib, pemigatinib, futibatinib) (Figure 6E-F, Supplemental Figure S7D-I). As expected, we observed minimal responses to FGFR inhibition in isogenic models that expressed the BRAF oncogenes. However, all four of the pan-FGFR inhibitors exhibited activity across our FGFR1-driven models, with IC50s in the nanomolar range (Figure 6E-F, Supplemental Figure S7A, S7C-H). Intriguingly, we also observed a spectrum of responses among FGFR1-altered lines, with the structural variants being the most sensitive and the SNV double mutant being the least sensitive (Figure 6E-F, Supplemental Figure S7A, S7C-H).

Our prior characterization of our model systems had revealed simultaneous activation of MAPK and mTOR pathway signaling in the absence of growth factor (Figure 3C-D). This is particularly relevant for FGFR1-driven gliomas that harbor co-mutations such as PTPN11, which can activate these pathways downstream of the FGFR1 receptor. Moreover, studies in other tumor contexts have revealed mTOR activation as a potential resistance mechanism for pan-FGFR inhibition^41,42^. We therefore hypothesized that combination approaches with agents to simultaneously inhibit MAPK or mTOR activity may enhance the efficacy of FGFR inhibitors.

We first tested this hypothesis *in vitro* using synergy assays across our FGFR1-altered mNSC lines with the FGFR inhibitor infigratinib in combination with either the MEK inhibitor trametinib or the mTOR inhibitor everolimus (Supplemental Figure S8A-B). BRAF V600E-driven mNSC models were included as a negative control. We observed no reproducible combinations of doses between trametinib and infigratinib that exhibited synergy across the FGFR1-altered lines (Supplemental Figure S8B). In contrast, combinations of infigratinib with the mTOR inhibitor everolimus exhibited synergy (Bliss synergy score >10) at low doses of each drug (Supplemental Figure S8A), but not in our negative control BRAF V600E model. These data support FGFR and mTOR inhibition as a potential combination therapy strategy in pLGGs.

### Diverse FGFR1 altered gliomas drive differential In vivo therapy effects of FGFR inhibitors

We next evaluated the efficacy of FGFR inhibition *in vivo*, leveraging the brain penetrant agent Infigratinib, both as single agent therapy, and in combination with either trametinib or everolimus. Dose tolerability studies were first performed to identify drug doses that were not toxic (Supplemental Figure S8C-D). Next, the striatum of SCID mice were injected with intracranial allografts of FGFR1::TACC1-expressing mNSCs and observed for glioma formation using MRI imaging, upon which mice were randomized to daily oral gavage treatment with either infigratinib (15mg/kg), everolimus (5mg/kg), trametinib (2mg/kg), infigratinib + trametinib (15mg/kg + 2mg/kg), or infigratinib + everolimus (15mg/kg + 5mg/kg) (Figure 6G).

The first cohort of mice with detectable gliomas initiated treatment on days 24 or 25 post injections, with a mean tumor volume as assessed by MRI of (5.65mm^3^ +/- 5.9mm^3^) (Figure 6H). Within this cohort, treatment of mice with FGFR1::TACC1 gliomas with infigratinib (either as single agent or in combination) significantly prolonged survival compared to mice treated with either trametinib or everolimus alone (*p*: 0.025, *p*: 0.025, *p*: 0.039, respectively) (Figure 6I). The median survival of mice treated with infigratinib was 16 days post initiation of treatment, compared to 15 or 16.5 days with everolimus or trametinib alone. However, no effects of combination therapy were seen compared to Infigratinib alone (Figure 6I).

The second cohort of mice with detectable gliomas initiated treatment six days later. However, these mice harbored significantly larger gliomas compared to those treated in the first cohort (5.65mm3 vs 21.1mm3, p value <0.0001) (Figure 6H), even though we had not detected gliomas by MRI imaging at the same time as Cohort 1. In this context, and with rapidly growing gliomas, we did not observe any survival advantage in any treatment arm, including with infigratinib. The median survival post initiation of treatment for mice treated with infigratinib within Cohort 2 was only 10 days (Figure 6J).

Together, these findings suggest a potential therapeutic benefit for single agent FGFR-inhibition in FGFR1::TACC1 expressing gliomas, however, with a potential window of opportunity with respects to glioma size and aggressiveness of growth.

### Human FGFR1-driven pLGGs exhibit moderate responses to the current generation of MEK and FGFR inhibitors

There are currently no FDA-approved treatments nor open clinical trials for children with FGFR-driven gliomas, and the optimal therapeutic approach remains unclear. This is particularly relevant in the setting of current MAPK-pathway inhibitors that have entered the clinical arena for pLGGs. We thus sought to perform a multi-institute retrospective analysis of patients diagnosed with FGFR-driven pLGGs, focusing on responses to standard chemotherapy, or following with MEK or FGFR inhibitors. This analysis included both unpublished cases integrated with a meta-analysis of published reports.

First, we performed a retrospective review of the 15 children with FGFR1-driven pLGGs diagnosed at DFCI that were included in our initial genomics analysis. Of these 15 patients, at a median follow up of 5.86 years, six children required further treatment with systemic chemotherapy (five with carboplatin/vincristine and one patient with single-agent carboplatin) (Figure 7A). One patient had a partial response to therapy while four patients had a best response of stable disease. The remaining patient was not evaluable (Figure 7A, Supplemental Table S5). The remaining nine children that did not require further treatment, with a median follow up of 5.16 years.

**Figure 7.**
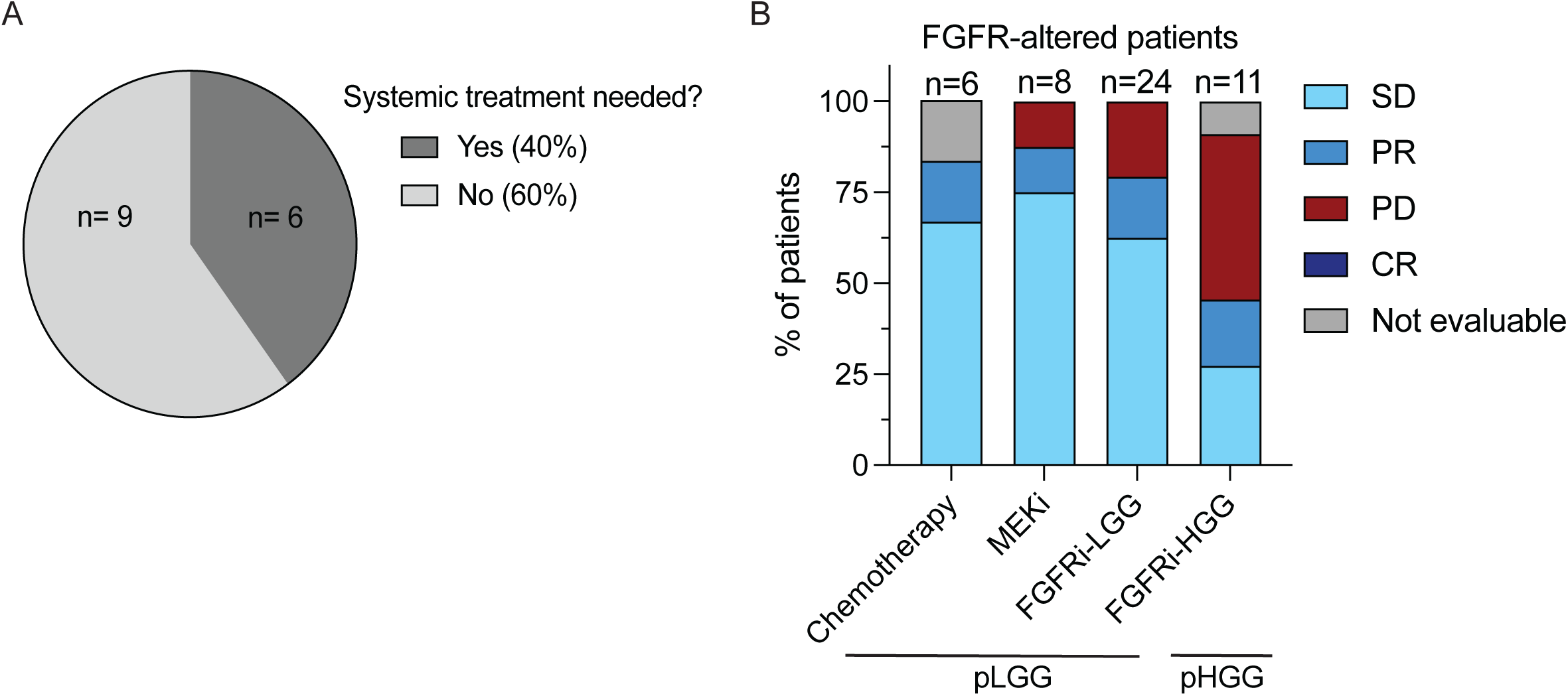
Treatment of FGFR1-altered gliomas with chemotherapy or targeted inhibitors show similar response rates. A) Pie chart showing the number of patients in FGFR-altered gliomas within the DFCI cohort that did or did not require further treatment following surgical management. B) Bar graph depicts the % of FGFR1-altered pediatric glioma patients treated with chemotherapy or targeted inhibitors with either FGFR- or MEK-inhibition, broken down by low-grade (pLGG) or high-grade (pHGG) histology. Chemotherapy-treated patients were from the DFCI cohort, targeted inhibitor-treated patients were from the DFCI cohort, published literature, and multi-institutional case studies. SD= stable disease, PR= partial response, PD: progressive disease, CR: complete response.

Across the larger meta-analysis, 43 episodes of targeted inhibitor treatment in 41 patients (32 patients with pLGGs) were identified, including patients treated with the pan-FGFR inhibitors Erdafitinib^43–46^ or Debio1347^47^, or MEK inhibition^48–51^. Two of these patients were treated with both a FGFR and MEK inhibitor as single agents at different timepoints. Of the 24 patients with pLGGs treated with FGFR inhibition, four children had a partial response, five had progressive disease while the remaining 15 patients had stable disease documented as their best response (Figure 7B). Six of the eight patients with FGFR-driven pLGGs treated with MEK inhibition had stable disease as their best response, with progressive disease and a partial response documented in the seventh and eighth patients (Figure 7B).

Overall, combining with patients treated with high-grade gliomas, the observed responses to targeted (FGFR or MEK) inhibitor therapy in pediatric FGFR-altered gliomas have been modest. Of 43 treatment courses identified, there were seven cases (16.2%) with partial responses, 24 (55.8%) with stable disease and 11 (25.6%) with progressive disease documented as the best response observed.

## Discussion

FGFR alterations are common across gliomas, particularly in pediatric patients. We find FGFR1 to be the most frequently altered FGFR family member in pediatric gliomas and highlight the heterogeneity of somatic drivers affecting them. Importantly, while we show glioma-associated FGFR1 alterations to be sufficient to drive gliomagenesis, we also highlight the need to optimize the efficacy of agents that target FGFR kinases themselves, or downstream activation of the MAPK and/or mTOR signaling pathways. While these FGFR driver events present the promise of precision medicine approaches, the path forward to the most effective strategies to target FGFR1 alterations remains elusive, even with the clinical development of pan-FGFR inhibitors.

Pediatric LGGs of different histological and molecular subtypes are treated with the same chemotherapy protocols, most commonly including vincristine and carboplatin. However, it is well documented that these gliomas exhibit heterogeneity of driver events, and varying underlying epigenetic and transcriptional signatures. Our findings clearly support FGFR-driven gliomas as being distinct from those driven by BRAF-alterations. Within the clinical cohort, FGFR1 alterations are enriched in glioneuronal pLGG types^5,21,22,24,52,53^, suggesting a potential link between FGFR1 expression and neuronal programs. This is further supported by our finding that overexpression of the FGFR1 alterations promote more neuronal transcriptional states compared to the BRAF alterations.

The therapeutic implications of these difference in underlying cell state are not well understood. For example, it has been reported that MEK inhibition in glioblastomas is associated with increased neuronal differentiation^54^. An alternate hypothesis is that MAPK pathway inhibition selects for pre-existing glioneural cell states that harbor primary resistance. Indeed, our isogenic FGFR1 NSC models exhibit less sensitivity to MAPK pathway inhibitors compared to those that harbor BRAF alterations. Further work is necessary to delineate these mechanisms which will be essential to inform strategies to optimize the use of these inhibitors in the clinical setting.

Generation of primary pLGG cell line models has been a major challenge for our field with the inability to propagate glioma cells *in vitro* or as patient-derived allografts^55^. Those that have been published have BRAF alterations, which excludes many of the other alterations, particularly FGFR1. To overcome this, we have generated multiple novel isogenic models of both mouse and human neural stem cells that are driven by FGFR1 and BRAF alterations. While they have the limitation that they are not patient-derived models, our isogenic systems provide a powerful tool to study each driver alteration and therapeutic approaches, both *in vitro* and as brain allografts in immunodeficient mice. While the intracranial gliomas grow as ‘high-grade’ tumors in the immune-deficient context, they do also appear to recapitulate some key features of the human gliomas. A particularly intriguing finding is the increased rate of intracranial hemorrhage that we observed in some of our FGFR1-altered and BRAF fusion gliomas, which has also been reported in the setting of children with pLGGs^56,57^. Further work is required to validate this initial observation, and to determine the mechanism through which FGFR1-altered gliomas increase the risk of hemorrhage.

We confirm FGFR1 alterations in pLGGs to exhibit heterogeneity. Both structural variants and somatic nucleotide variants are sufficient to generate activating driver events across pediatric gliomas. While structural variants involving FGFR proteins frequently occur as single driver events, similar to the majority of pLGGs overall, two hotspot SNV mutations (N546K and K565E) commonly co-occur with other mutations in FGFR1 itself, or in addition to other genes that activate MAPK and/or mTOR signaling (for example mutations in PTPN11 and PI3K pathway members). Intriguingly, we observed differences in the patterns of co-occurring alterations. FGFR1 N546K mutant gliomas harbor additional mutations that occur primarily in the extracellular domain, while FGFR1 K656E mutant gliomas have FGFR1 co-mutations that exclusively co-occur in the kinase domain (Supplemental Figure S1D).

Patterns of co-occurring mutations within the extracellular domain or the FGFR1 kinase may reflect different processes through which the FGFR1 SNVs enhance gliomagenesis which will be important to dissect to inform optimal therapeutic approaches. For example, mutations in the extracellular domain may influence ligand binding and specificity, compared to kinase mutations that may further contribute to its aberrant activation. Finally, for those gliomas with FGFR1 SNVs that did not harbor second SNVs in FGFR1 itself, they frequently co-occurred with other alterations with known roles in activating the MAPK and/or mTOR pathways, including PTPN11, NF1 and PIK3CA. Our modelling of the PTPN11 SNV suggests that it is insufficient to transform NSCs or induce glioma formation as a single driver event. Further investigation into these patterns and the functional role of their additional mutations will be critical to understand drug treatment and resistance to targeted inhibitors.

Our finding that FGFR-driven gliomas also exhibit heterogeneity across locations in which they arise has important clinical implications, particularly related to accurate diagnosis. While FGFR2 and FGFR3-driven gliomas most commonly occur in the cerebral hemispheres, almost 20% of all FGFR1-driven tumors arise in the midline structures including the thalamus and brainstem. Given these tumors are frequently diagnosed through stereotactic biopsies that can yield smaller amounts of tissue, it is imperative that all sequencing panels used in their molecular work-up are sufficient to detect each of the FGFR1-driver events.

Finally, our study confirms the inherent therapeutic challenges associated with FGFR1-driven pLGGs. Despite being the second most frequent somatic driver alteration of pLGGs, the optimal therapeutic approach remains unknown. Our clinical cohort suggests modest responses to both conventional chemotherapy and treatment with MEK inhibitors, a finding that is partially reinforced by our model systems that exhibited less sensitivity to MAPK pathway inhibition in the setting of FGFR1 alterations. There are currently no clinical trials evaluating the efficacy of FGFR inhibitors for children with pLGGs. However, our meta-analysis of the use of some of these FGFR inhibitors in the clinical setting has also revealed modest responses, with stable disease often reported as the best response. While penetrance across the blood-brain-barrier likely contributes to the lack of responses, our *in vivo* studies using a brain-penetrant inhibitor was also associated with transient responses, and only in mice in which treatment was initiated with smaller glioma sizes. It is possible that systemic toxicity has required the use of suboptimal doses, both in our mouse experiments, and more broadly in the early phase clinical trials. Indeed, it is important to note that the use of FGFR inhibitors in some of these trials was associated with a significant rate of bone complications, including fractures^44^. Future work is essential to identify combination approaches to optimize the efficacy of FGFR inhibitors, and to allow dosing using schedules that minimizes long-term toxicities.

## Acknowledgments

The authors gratefully thank and acknowledge the following funding sources: The Dana-Farber Pediatric Low-Grade Glioma Program (PB, KLL, EF, ME, SB), The Everest Centre for Low-grade Paediatric Brain Tumours (GN-000707, The Brain Tumour Charity, UK; DS, OW, DJ), Team Kermit and the Jared Branfman Sunflowers for Life Fund (PB, KLL, SC), Team Brainstorms (PB, KLL), Pediatric Brain Tumor Foundation (PB, KLL, EF, ME), V Foundation (PB and KLL), Team Jack Foundation (PB, KLL), Jack’s Drive 55 (PB, KLL), Damon Runyon-St. Jude Foundation Fellowship (03-24) (AA), The Swedish Childhood Cancer Fund (EM), DFCI – BMS (KLL), NIH, MMRF (KLL), The Lady Davis Institute/Dr. Arthur Rosenberg Memorial Fellowship Graduate Scholarship Fund and the James O and Maria Meadows Award from the Faculty of Medicine and Health Sciences at McGill University (BC), the EVEREST program of the Brain Tumor Charity and from the Dietmar Hopp Foundation (KiTZ Clinical Trial Unit 2.0) (OW).

LOGGIC Core BioClinical DataBank is supported by The Everest Centre for Research into Paediatric Low-Grade Brain Tumours (GN-000707, The Brain Tumour Charity, UK) and the PLGA Fund at the Pediatric Brain Tumor Foundation. The MNP2.0 study was supported by the German Childhood Cancer Foundation (DKS 2015.01).

We thank the Dana-Farber/Harvard Cancer Center in Boston, MA, for the use of the Specialized Histopathology Core, which provided histology and immunohistochemistry service. We thank the Molecular Biology Core Facilities at the Dana-Farber Cancer Institute for QC and sequencing of bulk and single-cell RNA-seq samples.

Finally, we would like to thank and acknowledge the many children and families affected by pediatric gliomas for their generous contributions to this research.

## Contributions

EM, AAA, SB, SO, HJ, ESF, MJE, QDN, DTWJ, KLL, PB conceived and designed experiments. EM, AAA, GA, JD, SB, PCP, MMC, DN, CCB, SM, SO, AC, PR, AD, LB collected data. EM, AAA, DS, SB, BC, PCP, RJ, JV, SM, JJ, JC, SH, CK, QDN performed analysis. DS, BC, JV, KM, MM, LAA, SHR, CMVT, ECH, PS, FS, KKY, TR, SC, KW, SP, AP, VL, SR, TB, AAS, MT, NJ, OW, CK, SA, DTWJ, KLL contributed data. EM, AAA, KLL, PB wrote the manuscript. All authors reviewed, edited, and approved the manuscript.

## Methods

### DFCI Patient cohort information

This study, including process for obtaining informed consent, was approved by the Dana-Farber Cancer Institute Institutional Review Board. To assess the landscape of FGFR-altered gliomas, clinical data and variant tumor calls were obtained through our institutional Precision Medicine Program sequencing database which contained clinical and research data from 2,514 primary CNS neoplasms included in this cohort. FGFR-altered gliomas underwent pathology and cytogenetic review for confirmation of fusion events, identification of focal copy number events (<10MB) an absence of fusion and identification of recurrent mutations within the cohort (>2). The review process identified 87 tumors with likely pathogenic FGFR alterations, referred to as the FGFR cohort.

### Oncopanel Sequencing

The DFCI cohort underwent next-generation targeted exome sequencing (OncoPanel) of cancer-related genes was performed using Illumina-based methods as previously described^58^. As this study was a retrospective analysis, some variability between cases exists due to changes in the design of this assay over the course of several years, including expansion of targeted genes from 300 to 471, with additional intronic coverage. We categorized each *FGFR* alteration described in clinical reports based on the FGFR gene that was altered (*FGFR1,2,3,4*) and by the category of genomic alteration (fusion/rearrangement, copy number, single nucleotide variant within protein coding regions and likelihood to be pathogenic).

### Copy arrays

Array comparative genomic hybridization (aCGH) was performed on DNA isolated from formalin-fixed paraffin-embedded (FFPE) tissue on the DFCI cohort using either the ThermoFisher Oncoscan CNV assay or the Agilent 1 × 1 M aCGH array, according to the manufacturers’ direction 922719973). All microarrays were analyzed using the Nexus Copy Number Software Package (BioDiscovery, El Segundo, CA) via the FASST2 segmentation with a significant threshold of 1.0E-12.

### RNA fusion panel

The presence of gene fusions within the DFCI cohort was also assessed using the clinical MGH Snapshot Fusion RNA-based assay that utilizes next generation sequencing from anchored primers within the known gene fusion partner (FGFR1, 3) and was performed on 6 x 5um sections of FFPE tissue^59^.

### MNP2 (KiTZ) cohort

This cohort was previously described^20^, and includes 425 gliomas from patients < 22 years at the time of diagnosis between 2015 and 2019. Tumors were classified by a superfamily class prediction score of > 0.9 for any glioma superfamily, irrespective of histology or scores for DNA methylation class families, classes, or subclasses. DNA methylation classes were called used the following classifier (version 12.5): https://www.molecularneuropathology.org/mnp/ > Classifiers > Classifier Versions > Brain classifier version 12.5. Most analyses were focused on the 30 patients with FGFR alterations, but excludes copy number analysis.

### Foundation Medicine (FM) cohort

The Foundation Medicine cohort of 502 FGFR-altered patients was derived from a collection of solid tumor clinical cases for which comprehensive genomic profiling had been previously performed^3^. Selected cases were verified primary CNS tumors harboring alterations in FGFR1-4 known from literature or included in Catalogue Of Somatic Mutations In Cancer (COSMIC) repository as well as those with likely functional status.

### Cell lines

Mouse neural stem cells (mNSC) obtained from the subventricular zone of CD1 mice^27^ were cultured as neurospheres in ultra-low adherence vessels (Corning, NY, USA) in tumor stem media (TSM) (500 mL Neurobasal-A (#10888-022), 500 mL DMEM/F12 (#11330-032), 10 mL HEPES Buffer Solution 1M (#15630-080), 10 mL MEM sodium pyruvate solution 100 mM (#11360-070), 10 mL MEM Non-essential amino acid solution 10 mM (#11140-050), 10 mL GlutaMAX solution (#35050-061), 10 mL Penicillin/Streptomycin (#15140122) and 1X B-27 Supplement Minus Vitamin A (from 50X) (#12587-010) from Invitrogen (MA, USA). When indicated, TSM media was supplemented with 12 ng/mL human-EGF (#78006), 12 ng/mL human-bFGF (#78006) and 2 ug/mL heparin solution 0.2% (#07980) from Stemcell Technologies (Vancouver, Canada). Neurospheres were dissociated using Accutase (#00-4555-56, Invitrogen) every 3-4 days and reseeded as single cells suspension with a density of 80-100,000 cells/mL in ULA flasks/plates.

Commercially available H9-derived human neural stem cells (GIBCO, Catalog nos. N7800-100, N7800-200) were immortalized by hTERT virus transduction (CMV-hTERT-Zeo lentivirus: Amsbio, LVP1130-Zeo). They were cultured as an adherent layer on geltrex-coated (Thermo Fisher, A1413302) plates in the TSM media listed above and accutase was used to detach them from plates.

### Model generation

FGFR1 transcript NM_023110.3 was used as a template for the generation of all FGFR1 SVs and SNVs. Similarly, for PTPN11, NM_002834.5 and TACC1 CCDS6109.1 exon 7-13 were used for SNV and SV respectively. BRAF transcript NM_004333.6 and KIAA1549::BRAF junction exon 15 to 9 was used. FGFR1 N546K – C1638A, FGFR1-ITD, FGFR1::TACC1 and PTPN11 E69K constructs were synthesized into Gateway compatible entry clones by GenScript with the addition of HA and V5 tags as indicated in the results segment. For the FGFR1-ITD, the following linker was used gatcgcagcctgcgcagcctgtgcagcttttggaaa (Whole-genome sequencing). Using the Gateway LR Clonase II (Thermo Fisher, 11791020), constructs were cloned into pLX311 (FGFR1s, BRAF, KIAA1549::BRAF, GFP and HcRed) or pLX307 (PTPN11 and LacZ).

To generate lentivirus, HEK293T cells were transfected with lentiviral expression vectors (10ug) and packaging plasmids encoding VSVG and PSPAX2 using the lipofectamine 3000 transduction kit (Thermo Fisher, L3000015), media was replaced after 6 hours and cell supernatant collected at 24 for lentivirus concentration using the Lenti-X concentrator (PT4421-2) according to manufacturer’s protocol.

mNSCs were infected using a spinfection protocol (2000 rpm for 2 hrs at 30 degrees C). Cells were selected with 0.75ug/mL puromycin (pLX307) for 3 days or 4ug/mL blasticidin (pLX311) for 7-10 days. ihNSCs were infected and selected with 0.5ug/mL puromycin (pLX307) for 3 days or 2.5ug/mL blasticidin (pLX311) for 7-10 days.

### Cumulative doubling assays

For cumulative growth assays, cells were seeded a density of 50-66,000 cells/mL (ihNSCs) or 60-100,000 cells/mL (mNSCs) in the presence or absence of FGF and EGF growth factors. Cells were counted every 3-4 days and when possible, a density of viable cells equal to the initiation of the experiment was reseeded. Doublings were calculated with the formula log2(total viable cells/seeded viable cells) at each passage and the cumulative results displayed. These experiments were carried out for 14 days.

### Western blots/densitometry analysis

Cells for immunoblotting were lysed on ice for 20-45 minutes in RIPA buffer contain protease and phosphatase inhibitors. Lysates were centrifuged at 13,000 x g for 10 min at 4 °C. Supernatant was collected and protein concentration quantified using the Pierce 660 nm Protein Assay (Thermo Fisher, 22660). Equal amounts of protein for each sample were aliquoted and mixed with 4X LDS Sample loading buffer and 10X NuPAGE Sample Reducing Agent and heated to 70 °C for 10 minutes. Lysates were loaded and run on NuPAGE Bis-tris 4-12% or NuPAGE Tris-Acetate 3-8% gradient gels.

The iBlot transfer system (Life Technologies, IB24001) was used to transfer protein to a PVDF membrane. The membranes were blocked in Advanblock (R-03726-E10) for one hour at room temperature. Subsequently, membranes were incubated with primary antibodies (listed in Supplemental Table S1) overnight at 4°C. Membranes were washed 3 x 5 minutes with 1X TBST and incubated with secondary HRP conjugated species-specific antibodies at room temperature for 1 hour, washed again, and then developed using SuperSignal West Pico PLUS Chemiluminescent Substrate (Thermo Fisher, 34578). Image capture was performed using the Fujifilm LAS-3000 Imaging System. Densitometry analysis was performed using Adobe photoshop. Full blots attached as a supplemental file.

### In vivo studies

All animal studies were performed according to Dana-Farber Cancer Institute Institutional Animal Care and Use Committee (IACUC)-approved protocols and included and equal number of male and female mice.

#### Cell injections

The mNSC lines described in Supplemental Figure S7A were injected stereotactically into the right striatum of 6 week-old female and male NSG mice (NOD.Cg-Prkdcscid Il2rgtm1Wjl/SzJ, The Jackson Laboratory, Bar Harbor, ME). Due to the number of mice injected, for all of the studies the mice cells were implanted on subsequent days. Briefly, mice were anesthetized with 2% isoflurane mixed with medical air and placed on a stereotactic frame. The skull of the mouse was exposed through a small skin incision, and a small burr hole was made using a 25-gauge needle at the selected stereotactic coordinates. The cells (300,000 cells in 3 µL PBS) were loaded on a 33-gauge Hamilton syringe and injected slowly using the following coordinates: - From Bregma, 0 mm AP, −2 mm ML, −2.5 mm DV. Upon completing injection, the needle was left in place for another minute, then withdrawn slowly to help reduce cell reflux. After closing the scalp with suture and staple, mice were returned to their cages placed on a warming pad and visually monitored until full recovery. Mice were then checked daily for signs of distress, including seizures, ataxia, weight loss, and tremors, and euthanized as they developed neurological symptoms, including head tilt, seizures, sudden weight loss, loss of balance, and ataxia.

#### MRI

MRI measurement of tumor volume was performed at the debut of neurological symptoms or around 2 months into the trial if asymptomatic for the study in Figure 5. For the drug study in Figure 6, the first MRI was performed 21/22 days post cell injections and again every subsequent week after until study was completed. MRI was performed using a Bruker BioSpec 7T/30 cm USR horizontal bore Superconducting Magnet System (Bruker Corp.). This system provides a maximum gradient amplitude of 440 mT/m and slew rate of 3,440 T/m/s and uses a 23 mm ID birdcage volume radiofrequency (RF) coil for both RF excitation and receiving. Mice were anesthetized with 1.5% isoflurane mixed with 2 L/min air flow and positioned on the treatment table using the Bruker AutoPac with laser positioning. Body temperature of the mice was maintained at 37°C using a warm air fan while on the treatment table, and respiration and body temperature were monitored and regulated using the SAII (Sa Instruments) monitoring and gating system, model 1025T. T2 weighted images of the brain were obtained using a fast spin echo (RARE) sequence with fat suppression. The following parameters were used for image acquisition: repetition time (TR) = 6,000 ms, echo time (TE) = 36 ms, field of view (FOV) = 19.2 x 19.2mm2, matrix size = 192 x 192, spatial resolution = 100 x 100 mm2, slice thickness = 0.5 mm, number of slices = 36, rare factor = 16, number of averages = 8, and total acquisition time 7:30 min. Bruker Paravision 6.0.1 software was used for MRI data acquisition, and tumor volume was determined from MRI images processed using a semiautomatic segmentation analysis software (ClinicalVolumes).

#### Drug treatment

NSG mice (6 weeks old, male and female) without tumors were used for the 10-day drug tolerability studies. For trametinib (HY-10999, MCE), 1 and 2mg/kg were tested, and 2mg/kg was well tolerated. For everolimus (HY-10218, MCE), 5 and 15mg/kg were tested and only 5mg/kg was well tolerated. For infigratinib (HY-13311, MCE), 10, 15, 20, and 30mg/kg were tested. 15mg/kg was the only dose well tolerated alone and in combination with the other drugs (Supplemental S8C-D).

Mice bearing FGFR1::TACC1-driven tumors were treated with 1) Vehicle control (10% DMSO, 40% PEG300, 5% Tween-80, 45% Saline), 2) trametinib (2mg/kg), 3) everolimus (5mg/kg), infigratinib (15mg/kg), 4) trametinib + infigratinib (2mg/kg + 15mg/kg), or 5) everolimus + infigratinib (5mg/kg + 15mg/kg) once daily by oral gavage. Drugs made in the same solution as the vehicle control and were prepared fresh prior to treatment.

#### Brain collection, tissue processing, and staining

The mice were sacrificed when at least one of the endpoints is reached, including: 15% loss in body weight from peak weight, poor body condition (BCS 2), signs of the animal being in morbid condition and neurological symptoms. After sacrifice, the mouse brains were collected and preserved, either by fixation or flash freezing. For fixation murine brains were fixed in 10% formaldehyde for 24 hr, then transferred to 70% ETOH until processing. Whole brain coronal sections were placed into cassettes in 70% ETOH and subsequently embedded in paraffin for block generation. Tissues from each paraffin block were cut at 5 μm sections and routine H&E staining was performed to be evaluated by an expert pathologist and digital pathology. Immunohistochemistry was performed on the Leica Bond III automated staining platform using the Leica Biosystems Refine Detection Kit (Leica; DS9800). FFPE tissue sections were baked for 30 minutes at 60°C and deparaffinized (Leica AR9222) prior to staining. Primary antibodies (Supplemental Table S1) with a 30M citrate antigen retrieval (Leica ER1 AR9961) were incubated for 30 minutes, visualized via DAB, and counterstained with hematoxylin (Leica DS9800). The slides were rehydrated in graded alcohol and cover slipped using the HistoreCore Spectra CV mounting medium (Leica 3801733).

#### Digital Neuropathologic Analyses

All slides were scanned at 40x magnification using a GT Leica Aperio scanner. An expert neuropathologist confirmed the presence or absence of tumor formation. Tumors were categorized as circumscribed, infiltrative, or mixed based on their border delineation with the surrounding normal brain tissue, as observed from H&E staining by a pathologist. The entire tumor area was annotated as a region of interest for further analysis. AI-based tissue segmentation for vasculature and hemorrhage quantification was performed using DenseNet v2 on the HALO AI image analysis platform (v3.0, Indica Labs, Albuquerque, NM). AI-based quantification of cellularity and IHC marker positivity was carried out using U-Net on Visiopharm image analysis software (v2023.09, Hørsholm, Denmark) for nuclear segmentation with customized apps and specific thresholds. Heatmaps for positive DAB signals were used for the automated selection of hotspot regions with the highest marker positivity for quantification.

### Drug response curves

Cells were seeded at a density of 1,000 cells/well in three technical replicates in corning 96-well plate (Corning, 3917). The indicated drugs (Supplemental table S1) were serial diluted before being added to the plate. Cells were incubated for 72h before viability was assessed by Cell-titer Glo (G7573). Cell titer glo was added per well (1:1), plates were incubated for 10 minutes at room temperature before readout of luminescence signal on a SpectraMax M5 plate reader. All results were normalized to vehicle control.

### Synergy assays

Cells were seeded in 384-well plates (Corning, 3765) at 1000 cells/well followed by drug dispensing (doses in Supplemental Table S1) with the HP D300e Digital Dispenser. Cells were incubated for 72h before viability was assessed by Cell-titer Glo (G9242). Cell titer glo was added per well (1:1), plates were incubated for 10 minutes at room temperature before readout of luminescence signal on the Pherastar plate reader. All results were normalized to vehicle control.

### Bulk RNA-seq

#### mNSC lines

Bulk RNA was extracted from mNSC cells using the Qiagen RNeasy kit (74104) and submitted to the Molecular Biology Core Facilities at Dana-Farber Cancer Institute. Libraries were prepared using Roche Kapa mRNA HyperPrep strand specific sample preparation kits from 200ng of purified total RNA according to the manufacturer’s protocol on a Beckman Coulter Biomek i7. The finished dsDNA libraries were quantified by Qubit fluorometer and Agilent TapeStation 4200. Uniquely dual indexed libraries were pooled in an equimolar ratio and shallowly sequenced on an Illumina MiSeq to further evaluate library quality and pool balance. The final pool was sequenced on an Illumina NovaSeq 6000 targeting 40 million 150bp read pairs per library at the Dana-Farber Cancer Institute Molecular Biology Core Facilities.

Sequenced reads were aligned to the UCSC mm10 reference genome assembly and gene counts were quantified using STAR (v2.7.3a)^60^. Differential expression analysis was performed using the R package DESeq2 v1.22.1^61^. RNAseq analysis was performed using the VIPER snakemake pipeline^62^. Genes with a baseMean (expression) higher than 50, an absolute log2FoldChange higher than 1.5 and a Benjamini and Hochberg corrected p-value of less than 0.05 were considered significant.

#### GSEA with MsigDB

Significant genes were run through the Molecular Signatures Database^30^. Negative log10(FDR) was calculated for each term and results were plotted in R using the package ggbarplot.

### Single-cell RNA-seq

Single-cell RNA-seq was performed on the mNSC lines using the Chromium Next GEM Single Cell 3**ʹ** Reagent Kits v3.1 (Dual Index) kit as per manufacturer’s instructions. 10,000 cells were loaded on the 10X controller for each sample. Libraries were quantified by Tape station 4200 (Agilent) and submitted to the Molecular Biology Core Facilities at Dana-Farber Cancer Institute. Uniquely indexed libraries were pooled in an equimolar ratio and sequenced on a NovaSeq 6000 S2 flowcell targeting 4 billion reads.

The CellRanger (v6.1.2) pipeline was used to map the sequencing data to the mouse genome (mm10). We then used the Seurat library (4.3.0) to convert the expression data to Seurat objects and perform quality control. First, we filtered for genes that appeared in at least 3 cells. Consistent with our expectation from mouse models, the fraction of reads that corresponded to mitochondrial RNA was very low (0.32% +/- 0.29%), thus we did not need to filter out cells by mitochondrial content. We filtered out indexes with fewer than 8000 reads to ensure we picked up cells that have one unique index. After filtering of low-quality cells, we obtained 19,009 cells total across the 4 samples. After performing log-normalization (scale.factor = 10000), we simplified the data set by selecting for the 3000 most highly variable genes. We next performed unsupervised clustering and dimensional reduction via RunPCA, FindNeighbors (30 dimensions), FindClusters (resolution = 0.5), and RunUMAP (30 dimensions) all through the Seurat library to create a two-dimensional visualization of the expression data.

### Analyses using publicly available datasets

#### KiTZ patient tumor-RNA-seq analysis

RNA-sequencing count data from pLGGs harboring FGFR1 or BRAF alterations were obtained and analyzed as previously described^28^. Normalization (mean of ratios) of the data and differential gene expression analysis between the BRAF (n=130) and FGFR1 (n=16) tumors was performed using the R package DESeq2^61^ with the Wald test. Genes with a baseMean (expression) higher than 50, an absolute log2FoldChange higher than 1.5 and a Benjamini and Hochberg corrected p-value of less than 0.05 were considered significant.

#### Retrieval and processing of bulk RNA-sequencing datasets

Bulk RNA-seq from the Evo-devo project^32^ and the Brainspan project^33^ were used for gene expression across brain developmental stages, and GTEx v8^34^ was used for gene expression across adult brain regions.

For the Evo-devo project, raw counts were downloaded and processed using an in-house RNA-seq pipeline as follows. Adapter sequences and the first four nucleotides of each read were removed from the read sets using Trimmomatic^63^ (v0.32). Reads were scanned from the 5′ end and truncated when the average quality of a four-nucleotide sliding window fell below a threshold (phred33 < 30). Short reads after trimming (<30 base pairs) were discarded. High-quality reads were aligned to the reference genome hg19 (GRCh37) using STAR^60^ (v2.3.0e) using default parameters. Reads mapping to more than 10 locations (MAPQ < 1) were discarded from downstream analyses. Gene expression levels were estimated by quantifying reads uniquely mapped to exonic regions defined by ensGene annotation set from Ensembl (GRCh37, N=60,234 genes) using featureCounts^64^ (v1.4.4). Normalization (mean-of-ratios) and variance-stabilized transformation of the data were performed using DESeq^61^ (v1.14.1). Brainspan RPKM normalized expression and metadata tables were downloaded and processed as follows: donors with <3 samples were removed, the provided brain region annotations were grouped into 16 unified regions, and expression data was log-transformed. GTEx v8 (GTEX) TPM normalized expression and metadata tables were downloaded and used directly.

#### Retrieval and processing of single-cell RNA-sequencing datasets

Single-cell RNA-seq atlases of mouse developmental^37,38^ and adult^65^ brain, as well as human developmental^36^ and adult^66,67^ brain were used to query cell type specificity of expression. Datasets were retrieved with cell type labels provided by authors and gene detection rate (% of cells with expression > 0) per cell type was calculated for visualization purposes.

A single-cell RNA-seq atlas of human fetal organogenesis^35^ was used to confirm early expression of FGFR3 in human embryo. Raw gene expression matrix and cell type labels from authors were retrieved and processed using Seurat^68^ (v4.3.0) as follows. Expression values were scaled to 10,000 UMI per cell and log normalized, as well as z-scored gene-wise. Dimensionality reduction was performed using principal component analysis (PCA) applied to the top 2000 most variant genes. The first 10 principal components were used as input for projection to two dimensions, using uniform manifold approximation and projection (UMAP)^69^.

### Statistical analysis

Log-rank (Mantel-Cox) tests were performed to analyze survival analysis of animal experiments. Fisher’s exact tests were used for clinical cohort comparisons. *p* values of <0.05 were considered significant.

## Data availability and code

Bulk and single-cell RNA-seq data from mNSCs have been deposited in GEO under accession number GSE274098. Patient cohort information, RNA-sequencing data are in supplemental files.

