## Supplemental Figures for "A diverse landscape of FGFR alterations and co-mutations defines novel therapeutic strategies in pediatric low-grade gliomas"

**Supplemental Figure S1**

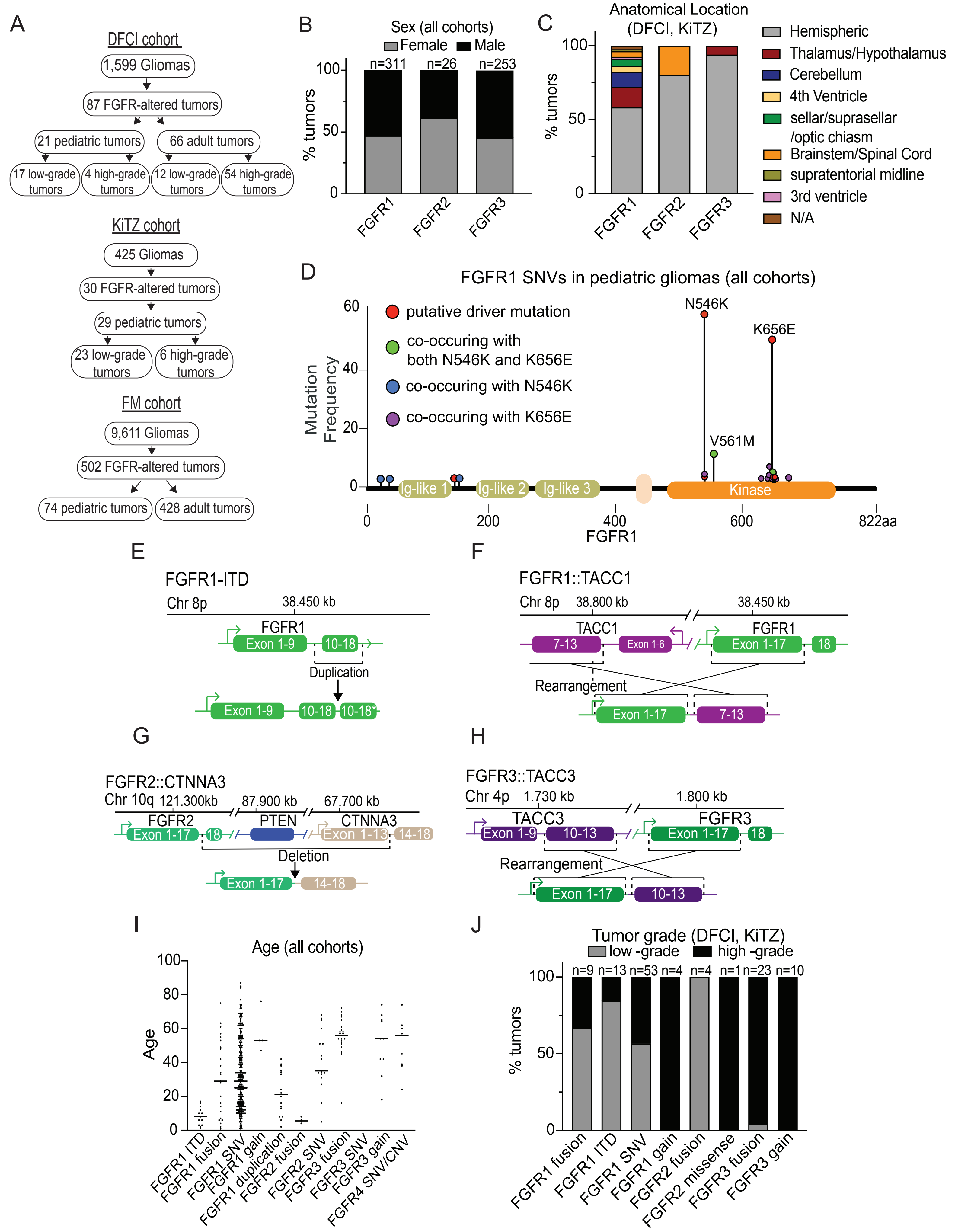


**Supplemental S1. Description of FGFR alterations in the three glioma cohorts**
A) Schematic summarizing the different glioma cohorts used to assess FGFR alterations. Distribution of FGFR gene alterations by B) patient sex and C) anatomical location. D) Lollipop plot showing the FGFR1 SNVs found in the three glioma cohorts. SNVs colored based on if they occur alone or co-occur with hotspot mutations. Schematics of the most frequently recurrent structural variants seen in the glioma cohorts including the E) FGFR1 kinase duplication (FGFR1-ITD), F) FGFR1::TACC1 fusion, G) FGFR2:CTNNA3 fusion, and H) FGFR3:TACC3 fusion. I) Age distribution across each specific FGFR alteration type in all glioma cohorts. J) Percentages of the tumor grade for each specific FGFR alteration type in the DFCI and KiTZ cohort. n= # of tumors in each group.

**Supplemental Figure S2**

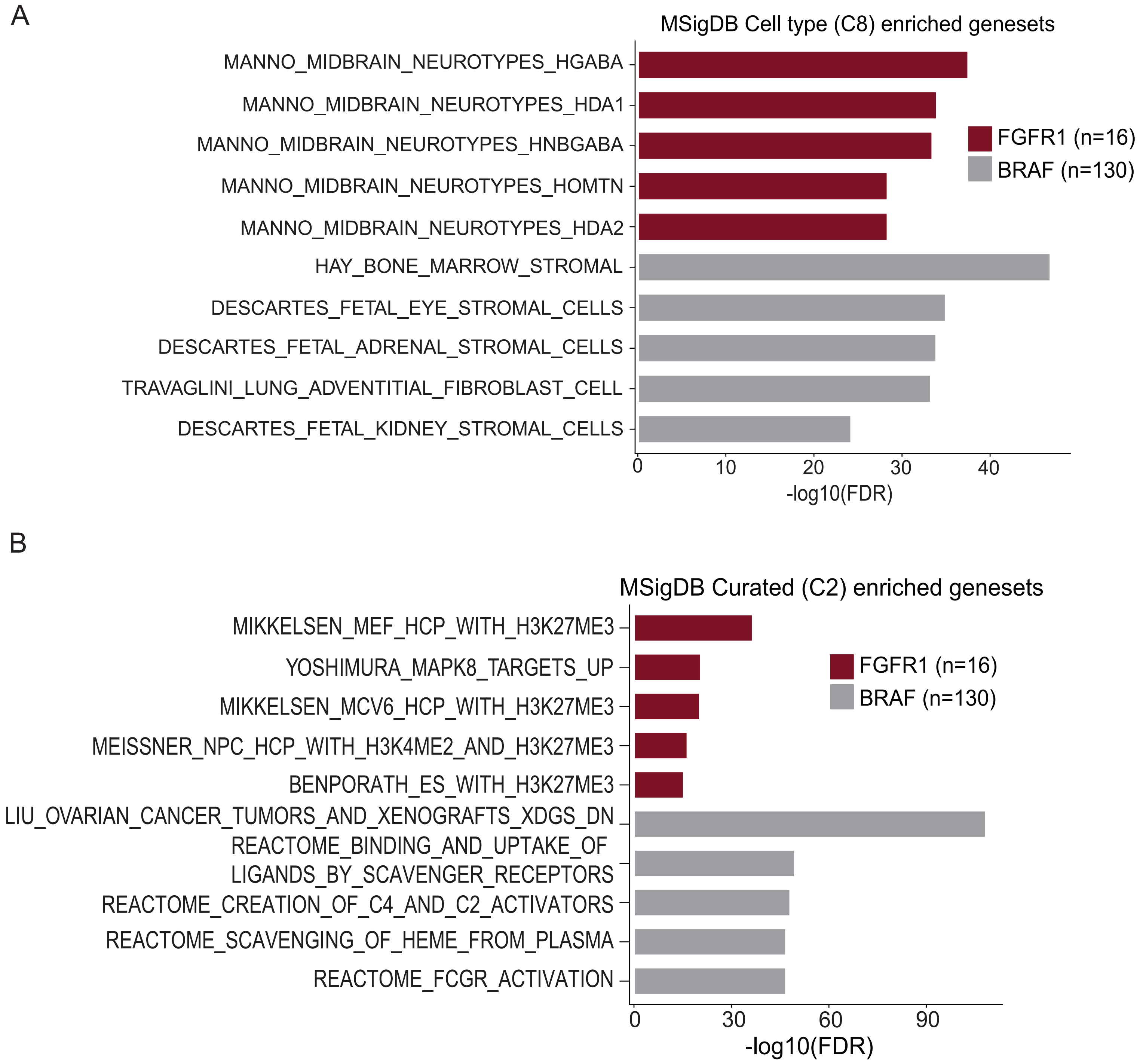


**Supplemental S2. FGFR1-altered tumors are enriched in neuronal cell type signatures**
A) Horizontal bar plots depicting the top five significant Cell Type C8 (MsigDB) terms enriched (ranked by significance) in FGFR1 (n=16) or BRAF (n=130)-altered patient tumors. B) Horizontal bar plots depicting the top five significant Curated C2 (MsigDB) terms enriched (ranked by significance) in FGFR1 (n=16) or BRAF (n=130)-altered patient tumors.

Supplemental Figure S3

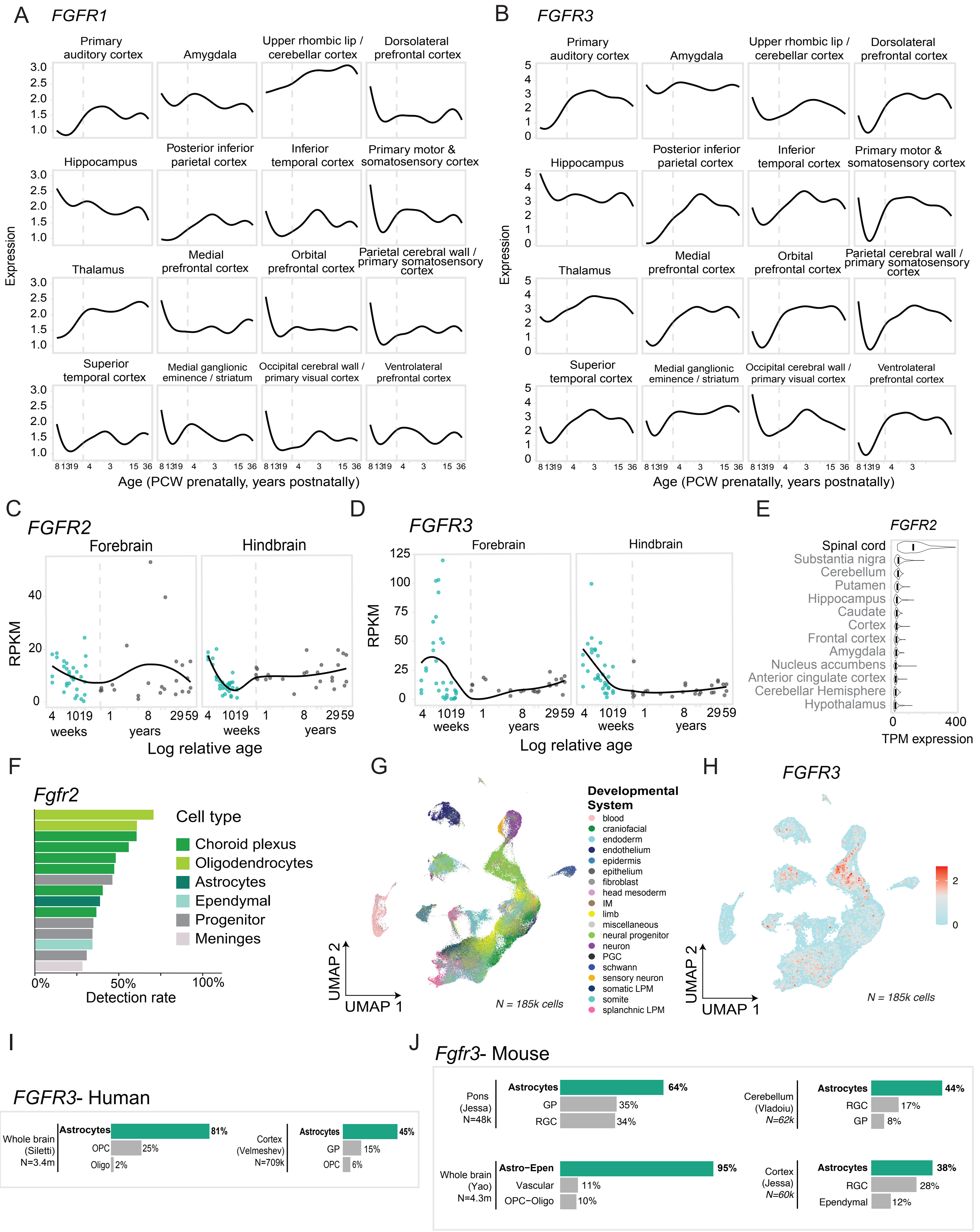


**Supplemental S3. Expression of FGFR family genes across developmental stages, cell types, and regions of the brain.**

A) Expression of FGFR1 across pre- and postnatal lifespan in a dataset of bulk RNA-seq samples. Each plot represents samples from an individual cortical or subcortical brain region. Vertical dashed line indicates birth. X axis denotes age and Y axis indicates normalized expression and are consistent across all plots. B) Expression of FGFR3 across lifespan as in panel A. C) Expression of FGFR2 in bulk RNA-seq across lifespan in human forebrain (left) and hindbrain (right). X-axis denotes sample age, measured in natural log of weeks post conception, and x-axis tick values correspond to weeks in prenatal points and years in post-natal points. D) Expression of FGFR3 in bulk RNA-seq across lifespan in human forebrain (n=55) (left) and hindbrain (n=59) (right). X-axis denotes sample age, measured in natural log of weeks post conception, and x-axis tick values correspond to weeks in prenatal points and years in post-natal points. Vertical dashed line represents birth. Y-axis depicts RPKM expression values. E) Expression (TPM) of FGFR2 in bulk RNA-seq samples across brain regions in adult human (GTEx v8). F) Top 15 cell clusters by detection rate (% of cells with expression of >0) of FGFR2 in developing mouse pons and forebrain. G) UMAP representation of human embryo 4-5.5 weeks post conception. Cells colored by cell annotation in the original study. H) UMAP representation of human embryo cells 4-5.5 weeks post conception. Cells colored by normalized expression of FGFR3. Color scale midpoint is set to the midpoint of maximum and minimum expression. I) Top three cell types by detection rate (% of cells with expression of >0) of FGFR3 in adult human brain. J) Top three cell types by detection rate (% of cells with expression of >0) of FGFR3 in developing mouse brain and adult mouse brain. Number of cells per dataset is indicated. OPC: oligodendrocyte precursor cells, Oligo: oligodendrocytes, GP: glial progenitors, RGC: radial glial cells.

### Supplemental Figure S4

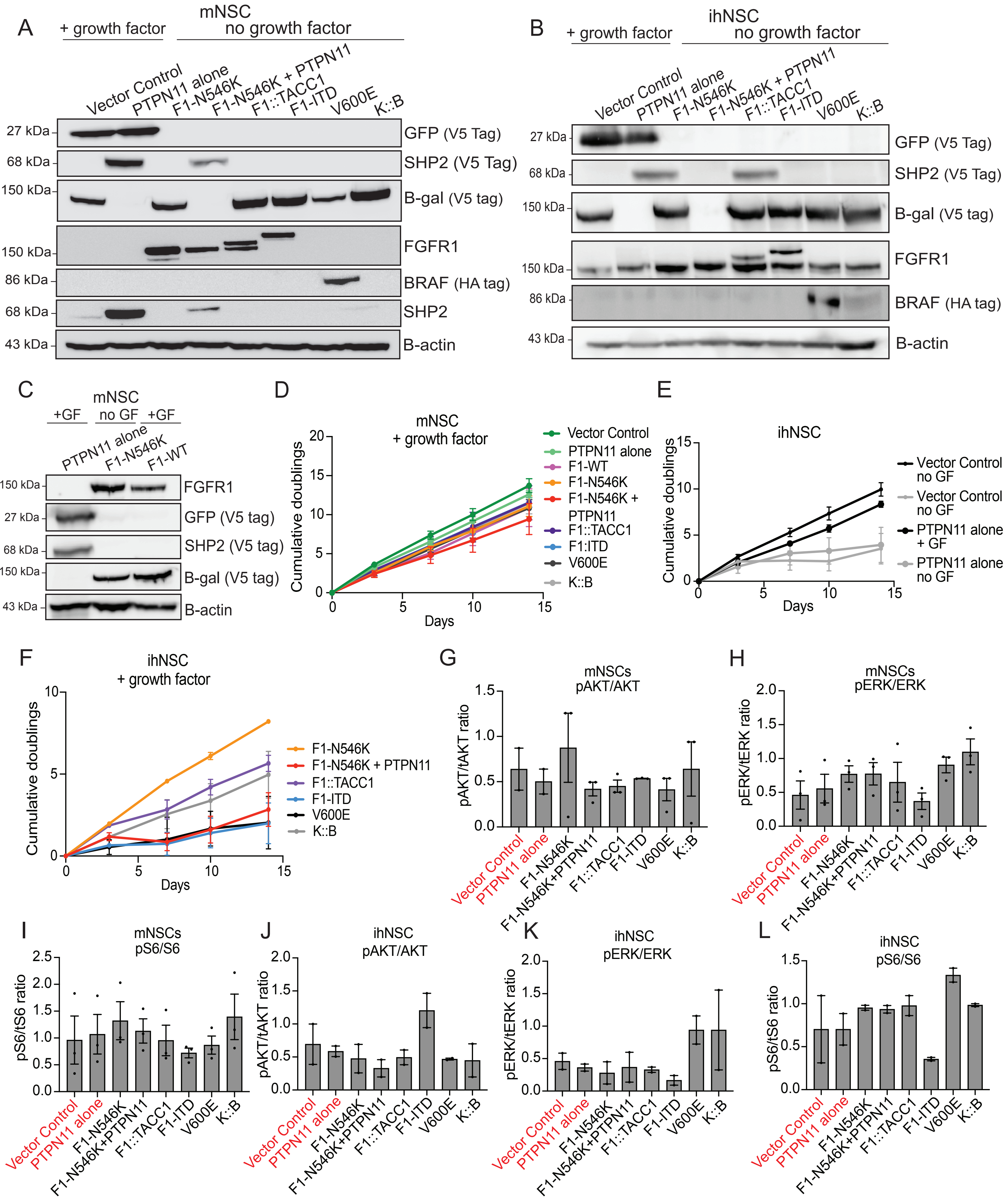


**Figure S4. Isogenic model validations**

Western blot of the overexpressed constructs in the A) mNSC and B) ihNSC isogenic lines. GFP, SHP2, and B-gal have V5 tags, while BRAF has an HA tag. ihNSCs have high endogenous FGFR1 expression. Vector Control and PTPN11 alone are grown in the presence of growth factor. C) Western blot validating the isogenic mNSC lines with FGFR1 WT overexpression. D) Cumulative doubling growth curves for the isogenic mNSC lines in the presence of exogenous growth factor. E) Cumulative doubling growth curves for the isogenic ihNSC Vector control and PTPN11 alone lines in the presence or absence of exogenous growth factor. F) Cumulative doubling growth curves for the isogenic ihNSC lines with FGFR1 or BRAF alterations in the presence of exogenous growth factor. Values and error bars represent the average +/- SEM of three independent experiments for cumulative doubling graphs. Densitometry quantification of G) pAKT/AKT expression, H) pERK/ERK expression, and I) pS6/S6 expression in the mNSC lines. Densitometry quantification of J) pAKT/AKT expression, K) pERK/ERK expression, and L) pS6/S6 expression in the mNSC lines. Vector Control and PTPN11 alone are grown in the presence of growth factor (red). GF= growth factor. For densitometry graphs, values and error bars represent the average +/- SEM of two or three independent experiments. Abbreviations for models= F1-N546K: FGFR1 N546K SNV, F1-N546K + PTPN11: FGFR1 N546K SNV + PTPN11 E69K SNV, F1-ITD: FGFR1-ITD, F1::TACC1: FGFR1::TACC1, V600E: BRAF V600E SNV, K::B: KIAA1549::BRAF.

### Supplemental Figure S5

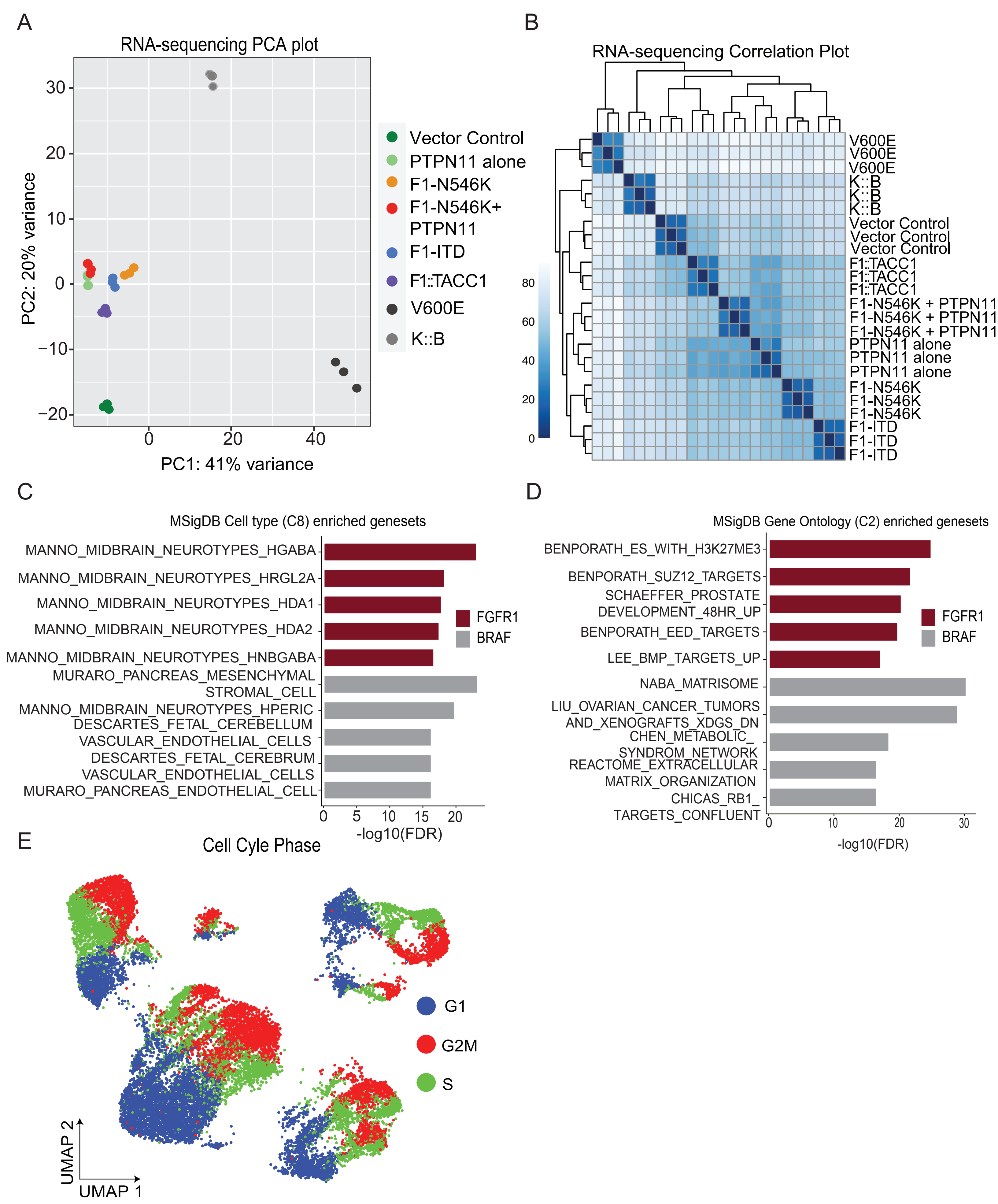


**Figure S5. RNA-seq analysis of mNSC lines**

A) PCA plot of bulk RNA-seq for isogenic mNSC lines, including three independent replicates per condition. B) Correlation heatmaps of the isogenic mNSC lines. C) Horizontal bar plots depicting the top five significant Cell type C8 (MsigDB) Terms enriched (ranked by significance) in FGFR1 (n=4) or BRAF (n=2)-altered mNSC lines. D) Horizontal bar plots depicting the top five significant Curated C2 (MsigDB) Terms enriched (ranked by significance) in FGFR1 (n=4) or BRAF (n=2)-altered mNSC lines. The x-axis depicts negative Log(FDR), showing significance of enrichment of each gene set. Labels indicate genes expressed by each cell line, Abbreviations for models= F1-N546K: FGFR1 N546K SNV, F1-N546K + PTPN11: FGFR1 N546K SNV + PTPN11 E69K SNV, F1-ITD: FGFR1-ITD, F1::TACC1: FGFR1::TACC1, V600E: BRAF V600E SNV, K::B: KIAA1549::BRAF E) UMAP embedding of the FGFR1-altered mNSCs colored by the cell cycle phase.

### Supplemental Figure S6

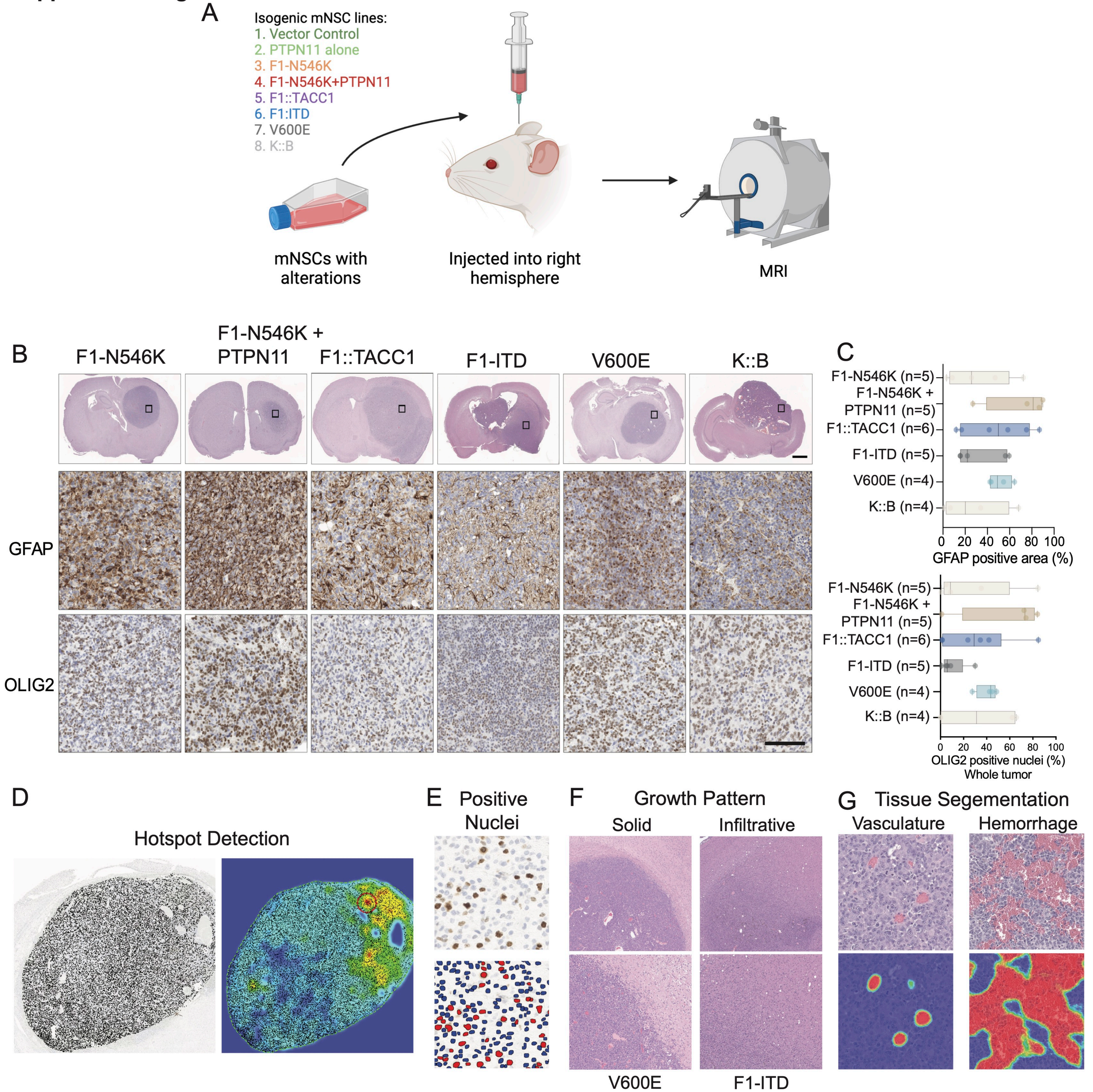


**Figure S6. Overview of digital pathology techniques**

A) Schematic of the in vivo study. 300,000 isogenic mNSC cells driven by the indicated alterations were injected into the right hemisphere of mice and tumors were assessed by MRI. B) Representative H&E images showing tumor formation at low magnification (1.5x, scale bar = 1 mm), with immunohistochemistry (IHC) demonstrating positivity for GFAP and OLIG2. Images of mouse brains are reshown from Figure 5 for context of location. C) AI- based digital quantification of GFAP and OLIG2 positivity in whole tumors using U-net on the Visiopharm platform. Illustrative images of histopathology characterization methods. # of mice is depicted. D) Heatmap demonstrating the hotspot region with the highest positivity for the DAB signal of a tumor stained with Ki-67 on the AI-based Visiopharm platform. E) Nuclei segmentation using U-Net on Visiopharm for positive nuclei quantification. F) Images showing a solid growth pattern of a circumscribed BRAF V600E tumor (left) and an infiltrative FGFR1-ITD tumor (right). G) Deep learning-based automated tissue segmentation of vasculature (left) and hemorrhage (right) using DenseNet on the HALO AI platform. Abbreviations for models= F1-N546K: FGFR1 N546K SNV, F1-N546K + PTPN11: FGFR1 N546K SNV + PTPN11 E69K SNV, F1-ITD: FGFR1-ITD, F1::TACC1: FGFR1::TACC1, V600E: BRAF V600E SNV, K::B: KIAA1549::BRAF.

**Supplemental Figure S7**

A IC50 doses

| | mNSC | | | | | | ihNSC | | | | | |
|---|---|---|---|---|---|---|---|---|---|---|---|---|
| | F1-N546K | F1-N546K +PTPN11 | F1::TACC1 | F1-ITD | V600E | K::B | F1-N546K | F1-N546K +PTPN11 | F1::TACC1 | F1-ITD | V600E | K::B |
| Tovorafenib | 702.1 nM | 466.4 nM | 3.592 uM | 3.167 uM | 116.9 nM | 319.9 nM | N/A | N/A | N/A | N/A | N/A | N/A |
| Belvarafanib | 56.57 nM | 52.49 nM | 155.9 nM | 155.5 nM | 25.01 nM | 8.92 nM | N/A | N/A | N/A | N/A | N/A | N/A |
| Trametinib | 1.02 nM | 0.71 nM | 3.32 nM | 3.28 nM | 0.087 nM | 0.156 nM | 354.1 nM | 48.61 nM | 258.2 nM | 699.1 nM | 3938 nM | 6028 nM |
| Everolimus | 4992 uM | 41.52 uM | 97.26 uM | 39.51 uM | 6.9 uM | 16.91 uM | 155162 uM | 43002 uM | 65233 uM | 144.7 uM | 163478 uM | 237744 uM |
| Infigratinib | 3.283 nM | 8.912 nM | 1.25 nM | 1.77 nM | 332.5 nM | 600.7 nM | 2.1 uM | 188.2 nM | 6.46 nM | 6.65 nM | 34455 uM | 5045 uM |
| Erdafitinib | 0.807 nM | 2.304 nM | 0.122 nM | 0.171 nM | 15.58 uM | 17.3 uM | 191.5 nM | 149.4 nM | 7.19 nM | 10.04 nM | 7.82 uM | 6.97 uM |
| Pemigatinib | 2.19 nM | 8.46 nM | 0.44 nM | 0.797 nM | 7.782 uM | 64246 uM | 118.6 nM | 240.5 nM | 4.139 nM | 8.37 nM | 6.89 uM | 3.6 uM |
| Futibatinib | 1.06 nM | 2.9 nM | 0.96 nM | 0.57 nM | 7388 uM | 154.4 uM | 15.73 nM | 23.58 nM | 2.017 nM | 2.177 nM | 51.47 uM | 49.91 uM |

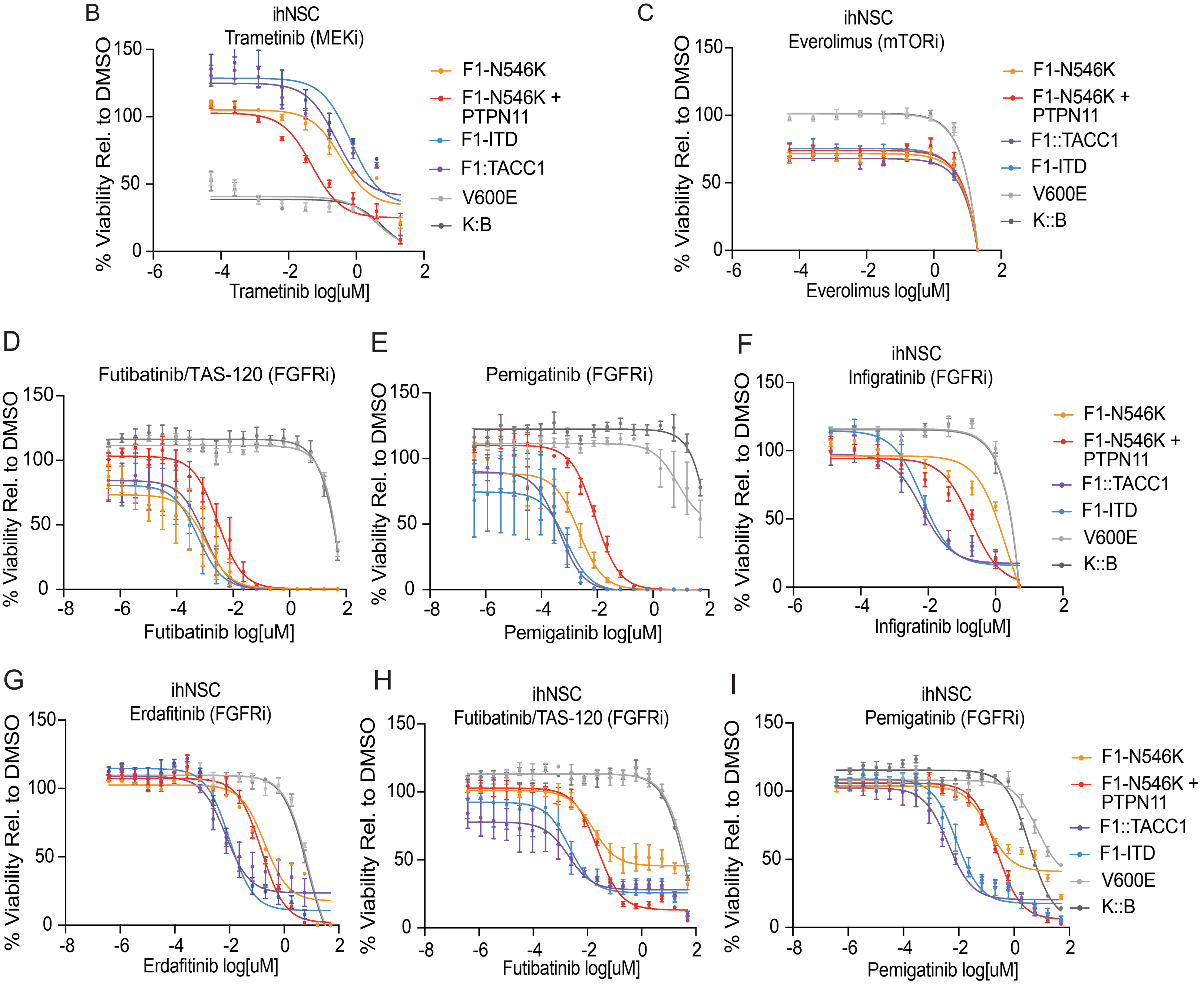


**Figure S7. FGFR1-driven models are sensitive to FGFR inhibition**

A) Table of IC50 values from dose response curves in the mNSC and ihNSC models. Dose response curves in the ihNSC models for B) trametinib (MEKi) and C) everolimus (mTORi). Dose response curves in the mNSC models for D) futibatinib (FGFRi) and E) pemigatinib (FGFRi). Dose response curves in the ihNSC models for F) infigratinib (FGFRi), G) erdafitinib (FGFRi), H) futibatinib (FGFRi) and I) pemigatinib (FGFRi). Values and error bars represent the average +/- SEM of three independent experiments. Abbreviations for models= F1-N546K: FGFR1 N546K SNV, F1-N546K + PTPN11: FGFR1 N546K SNV + PTPN11 E69K SNV, F1-ITD: FGFR1-ITD, F1::TACC1: FGFR1::TACC1, V600E: BRAF V600E SNV, K::B: KIAA1549::BRAF.

### Supplemental Figure S8

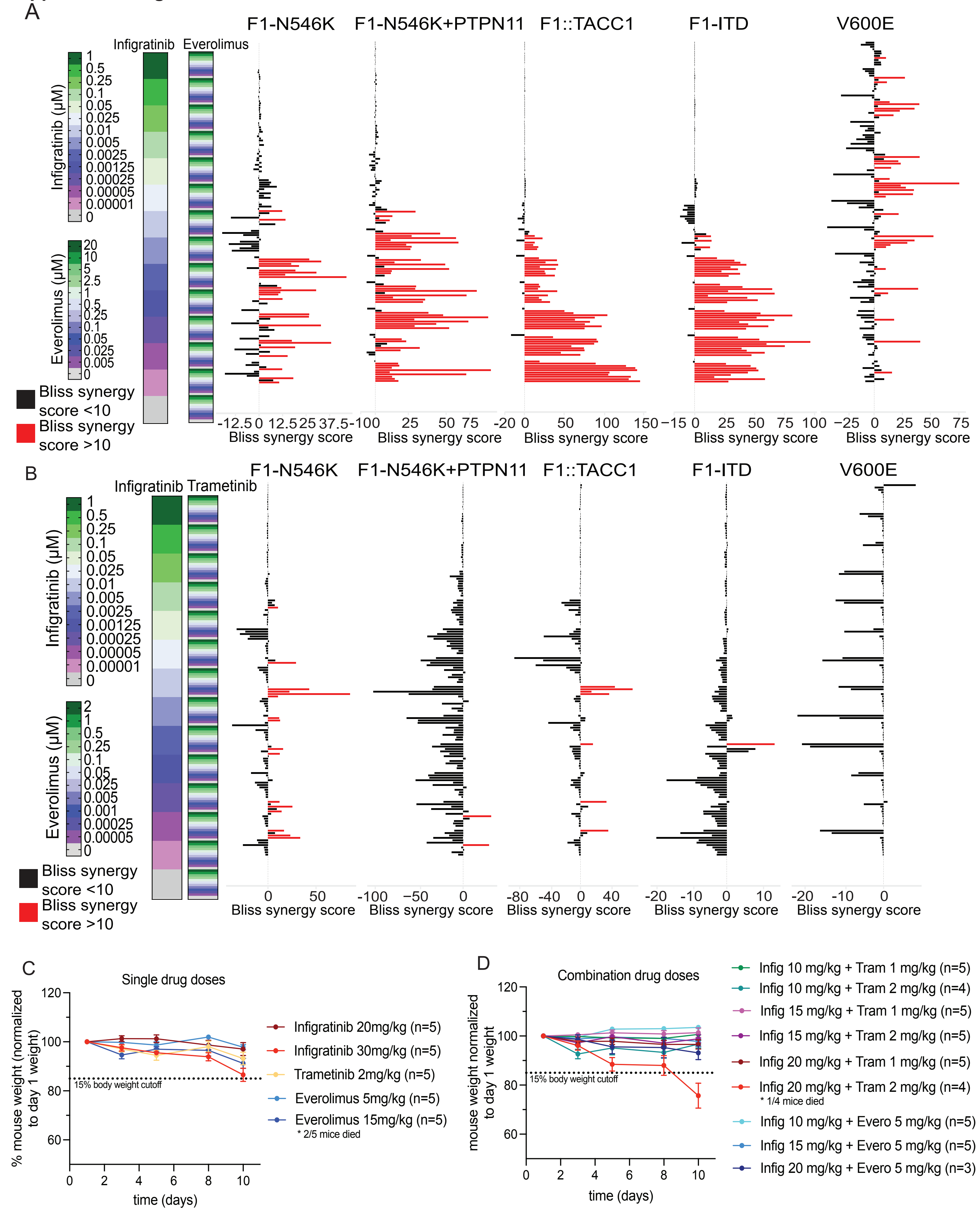


**Figure S8. Drug synergy plots and tolerability study**

A) Schematic summarizes drug synergy results for the combination of infigratinib and everolimus in FGFR1-altered and BRAF V600E mNSC lines. Infigratinib and everolimus combination doses are shown in the green to purple scale on the left of the plot. For each dose combination the bliss synergy score is plotted. Synergy is defined by a bliss score >10 (red bars). B) Schematic summarizes drug synergy results for the combination of infigratinib and trametinib in FGFR1-altered and BRAF V600E mNSC lines. Infigratinib and trametinib combination doses are shown in the green to purple scale on the left of the plot. For each dose combination the bliss synergy score is plotted. Synergy is defined by a bliss score >10 (red bars). Synergy experiments represent three technical replicates. Mouse weight in the drug tolerability study over 10 days of C) single or D) combination drug doses of infigratinib, everolimus, and trametinib. Abbreviations for models= F1-N546K: FGFR1 N546K SNV, F1-N546K + PTPN11: FGFR1 N546K SNV + PTPN11 E69K SNV, F1-ITD: FGFR1-ITD, F1::TACC1: FGFR1::TACC1, V600E: BRAF V600E SNV, K::B: KIAA1549::BRAF.
